# InGene: Finding influential genes from embeddings of nonlinear dimension reduction techniques

**DOI:** 10.1101/2023.06.19.545592

**Authors:** Chitrita Goswami, Namrata Bhattacharya, Debarka Sengupta

## Abstract

We introduce *InGene*, the first of its kind, fast and scalable non-linear, unsupervised method for analyzing single-cell RNA sequencing data (scRNA-seq). While non-linear dimensionality reduction techniques such as t-SNE and UMAP are effective at visualizing cellular sub-populations in low-dimensional space, they do not identify the specific genes that influence the transformation. *InGene* addresses this issue by assigning an importance score to each expressed gene based on its contribution to the construction of the low-dimensional map. *InGene* can provide insight into the cellular heterogeneity of scRNA-seq data and accurately identify genes associated with cell-type populations or diseases, as demonstrated in our analysis of scRNA-seq datasets.

## 1. Introduction

Dimensionality reduction is often used to visualize high dimensional and complex data. The ability to represent multi-dimensional samples on a two-dimensional scatterplot lends a multitude of benefits. It helps us investigate the patterns of similarity between individual samples and can also aid in discovering rare entities (1). When the dimensionality reduction technique is linear in nature, such as Principal Component Analysis (PCA), we can also decipher which features contribute to the variability of the data and thus hold higher importance. Interpreting feature importance is essential for understanding the data, model analysis and improvement. When it comes to biological research, it is exceptionally crucial that one can analyze and interpret both samples and their features (e.g. genes).

The rapid progress in next-generation sequencing (NGS) in recent years has provided many valuable insights into complex biological systems. In addition to revealing an incredible amount of information about fundamental biology and disease, high-throughput sequencing technologies are becoming increasingly versatile. They are applied to various clinical problems, ranging from cancer genomics to diverse microbial communities. NGS-based technologies for genomics, transcriptomics, and epigenomics are now increasingly focused on the characterization of individual cells. Several studies have shown that genomic alterations are more heterogeneous at the single-cell level than at the bulk level (2).

The usual number of cells analyzed in a single-cell study ranges between a few hundred to several thousand (3). Dimension reduction techniques are routinely used to facilitate downstream analyses and visualization of such high dimensional datasets. Such methods replace several thousands of genes with a small number of latent variables while preserving important patterns in datasets. In the field of single-cell transcriptomics, Principal Component Analysis (PCA) (4), t-Distributed Stochastic Neighbor Embedding (t-SNE) (5) and Uniform Manifold Approximation Projection (UMAP) (6) are used the most (7; 8). t-SNE and UMAP both claim to position similar cells closer together. Becht et al. (9) argued that UMAP is preferable to t-SNE because it better preserves the global structure of the data and is more consistent across runs. That being said, both tSNE and UMAP are immensely popular tools today for data visualization. PCA uses a linear transformation to project individual cells into a latent space while maximizing the variance of the projections. On the contrary, t-SNE and UMAP identify hidden structures in data, exposing natural clusters and non-linear variations along the dimensions (5) . Following dimension reduction, an obvious question to ask is, “What are the key genes?”. For PCA, often genes with the highest loadings in the first few principal components are considered to be the most influential (10; 7). t-SNE and UMAP are non-linear dimension reduction techniques that can identify hidden data structures and expose natural clusters and non-linear variations along the dimensions. These techniques are often used to visualize high-dimensional data, but they do not have an equivalent technique to identify the most influential genes as PCA does with loadings. We propose *InGene*, a machine learning based tool for identifying genes that best explain non-linear transformations for single-cell transcriptome data.

## 2. Results

### 2.1. Overview of *InGene*

Popular feature selection techniques to detect highly variable genes are generally dispersion based methods such as -Coefficient of variation (CV) (11), Fano Factor (12) and Gini Coefficient or Gini Index (13). They are all univariate and are calculated using the mean and standard deviation of the features. The drawback of rank ordering genes using these techniques are quite a few. Using a variance based method means that low level expression and low variability features will be under-sampled due to their low variability values. In addition, genes with higher expressions and variability values have a higher probability of showing up. Also, such methods do not capture the relative difference in expression values. For example, genes with high expressions and low variability in cells may not appear believable since their relative difference in expression value will be close to zero compared to low expression and high variability cells. Please note that dispersion-based measures of heterogeneity are often overemphasized, which can result in the selection of marker genes that are actually weakly associated with the true sub-population.

To overcome this, we believe that the cellular arrangement of the sub-population should be considered while determining essential marker genes. Thus, *InGene* is designed to utilize the structure of the sub-population during the selection of marker genes.

Dispersion based methods also do not do a good job of conveying the inherent relationships between the variance in transcript levels between cells and the variables believed to be essential for separating cell types. The methods can not fully utilize the biological knowledge of the data structure, such as co-expression or similarity structure. Therefore, methods that focus on highly variable genes based on the scores given by dispersion methods can be inadequate. To circumvent the above issues, we skipped selecting only the highly variable genes and instead chose to retain all the genes that are expressed post gene filtering in *InGene*.

Another popular technique to rank order genes is Principal Component Analysis (PCA) (14). PCA is a dimension reduction method of transforming high dimensional data into low dimensional space. PCA provides an informative basis for attribute ranking or exploring the correlation between the genes. Though it circumvents some of the drawbacks of dispersion based methods, PCA is not very efficient when it comes to preserving local structures. Being a linear method, it also fails to perform adequately for more heterogeneous datasets.

Non-linear dimensionality reduction methods, such as t-distributed stochastic neighbor embedding (tSNE) (5) and uniform manifold approximation and projection (UMAP) (6) are used to construct an embedding of the data that preserves pairwise distances across the projected low-dimensional space. These methods are designed for the embedding of high dimensional data into some of its low dimensional projections. They can be used to obtain low dimensional representations of a set of data that preserve the relationships among high-dimensional projections are such as pairwise spatial distribution. Though much more potent in retaining the structure of the data in low dimensional space when compared to PCA and dispersion methods, non-linear dimensionality reduction methods have remained only as a mere visualization tool. Due to their non-linear nature, one is unable to decode which genes carry a heavier weightage (importance) in the construction of the lower dimensional embeddings. Thus these techniques remain underutilized when it comes to analysis. While visual interpretation does help one decipher the patterns in a dataset, it is pertinent to decode the key features influencing those patterns, especially when dealing with biological and clinical datasets.

Figure 1 describes how *InGene* extracts the features that best explain the cellular arrangement in the latent dimension (see Methods).

**Fig. 1:**
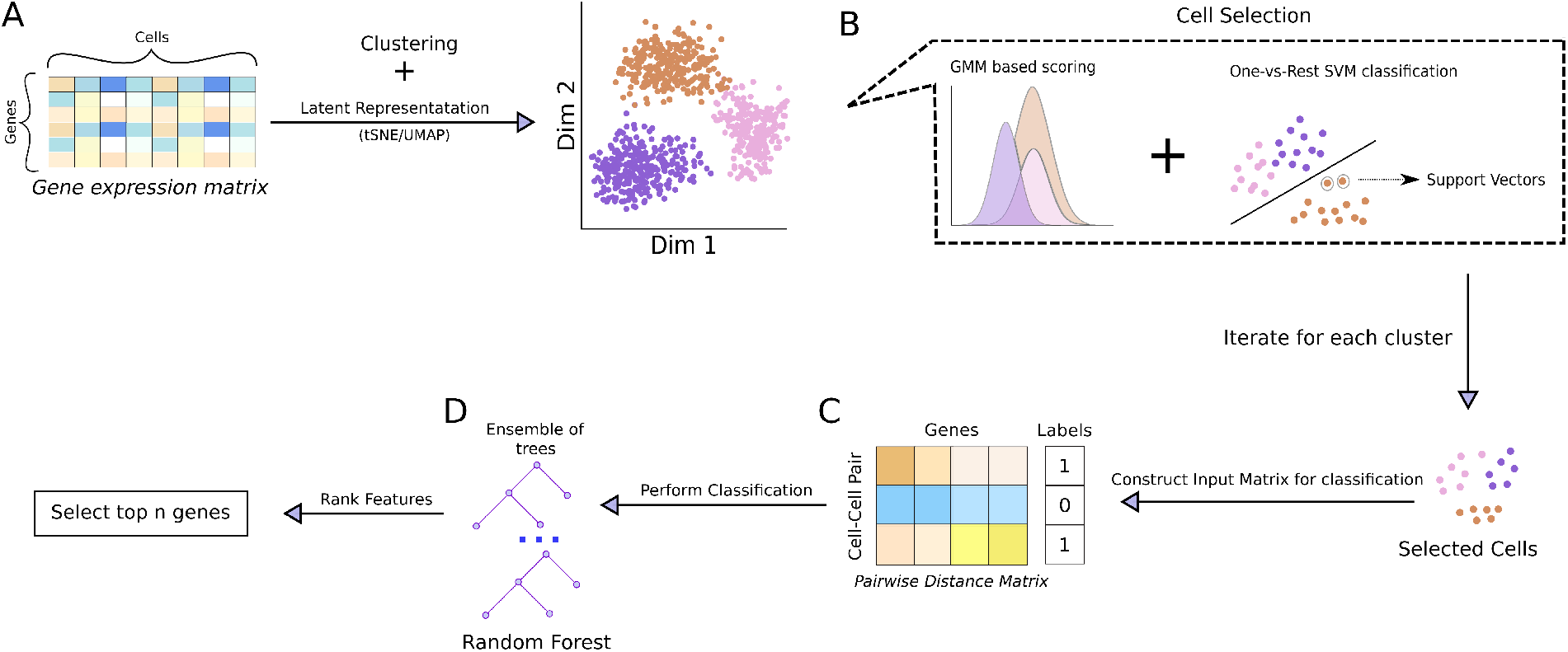
Schematic representation of workflow. **(A)** The gene expression data is clustered (using Seurat pipeline). The results are projected onto a 2D latent representation using t-SNE/UMAP. **(B)** Cells are selected from each cluster using the latent representation. The subsampling is done using one-vs-rest SVM, combined with likelihood values from GMM. (C) Subsampled cells are combined pairwise to construct the feature matrix to be used for feature ranking. Each pair of cell is a new entry, and labeled as ‘1’ if both the cells belong to same cluster or ‘0’ otherwise. **(D)** The feature matrix is passed to Random Forest classifier, which also ranks the features (genes).

*InGene* was compared with popular unsupervised and supervised methods that rank orders genes for single-cell transcriptomic data - CV^2^ (11), Fano Factor (12), Gini Coefficient or Gini Index (13), PCA (4), MAST (15), scGeneFit (16). The same is evident when we compare the reconstructed tSNE/UMAP using only the top-ranked *InGene* genes (max 500) along with the one that was constructed using all the genes.

The observation was consistent for all the scRNA-seq datasets we applied *InGene* on : Melanoma(Figure 2, Figure 3), PBMC 68K(Figure S3,S4,S5,S6), Darmanis(Figure S7, S8, S9, S10). Each dataset has UMAP and tSNE representation for both the ground truth annotations and the clustered annotation.

**Fig. 2:**
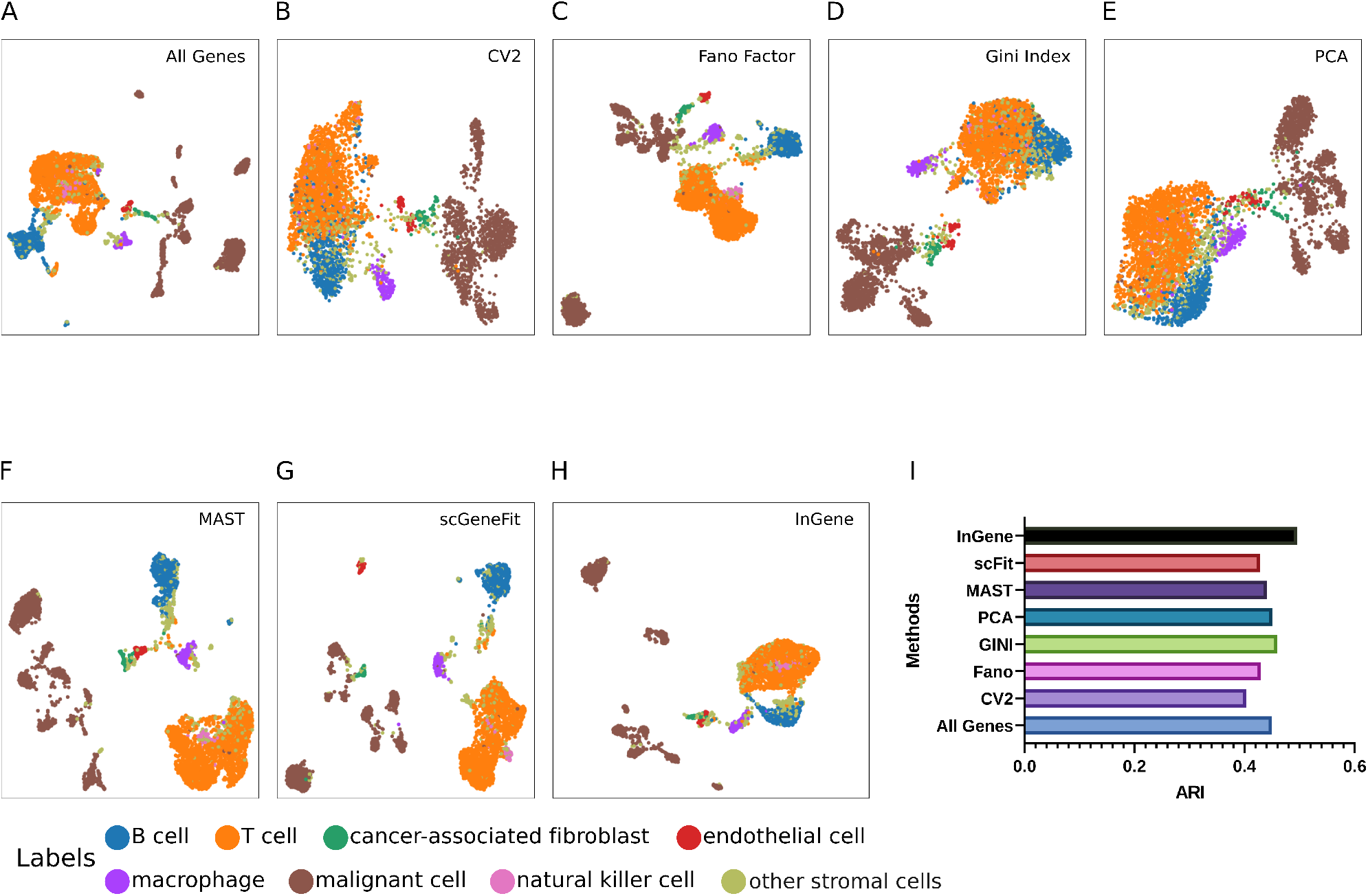
Explaining UMAP for single-cell melanoma dataset with *InGene*. **(A)** UMAP constructed with all the genes post-filtering. **(B)** UMAP constructed with top 500 CV2 genes. **(C)** UMAP constructed with top 500 Fano Factor genes. (D) UMAP constructed with top 500 Gini Index genes. **(E)** UMAP constructed with top 500 PCA genes. **(F)** UMAP constructed with top 500 MAST genes. **(G)** UMAP constructed with top 500 scGeneFit genes **(H)** UMAP constructed with top 500 *InGene* Factor genes. **(I)** ARI scores for the different methods. The gene set from each methods is used to cluster the dataset, using Leiden algorithm. The cluster labels obtained are then compared with the true labels to obtain the ARI values.

**Fig. 3:**
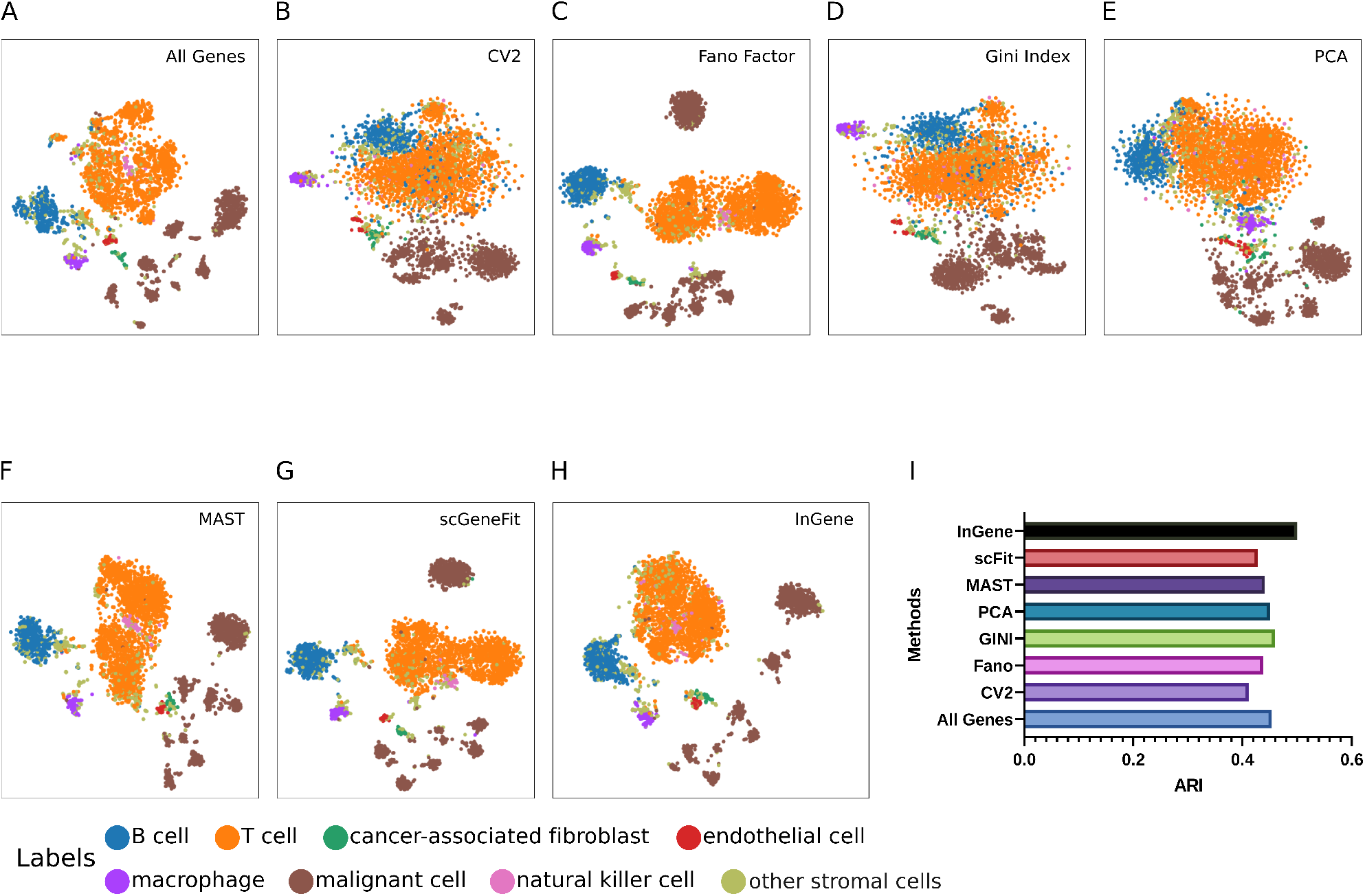
Explaining t-SNE for single-cell melanoma dataset with *InGene*. **(A)** t-SNE constructed with all the genes post-filtering. **(B)** t-SNE constructed with top 500 CV2 genes. **(C)** t-SNE constructed with top 500 Fano Factor genes. **(D)** t-SNE constructed with top 500 Gini Index genes. **(E)** t-SNE constructed with top 500 PCA genes. **(F)** t-SNE constructed with top 500 MAST genes. **(G)** t-SNE constructed with top 500 scGeneFit genes **(H)** t-SNE constructed with top 500 *InGene* Factor genes. **(I)** ARI scores for the different methods. The gene set from each methods is used to cluster the dataset, using Leiden algorithm. The cluster labels obtained are then compared with the true labels to obtain the ARI values.

### 2.2. *InGene* reveals relevant genes

*InGene* was evaluated on four datasets comprising various diseases, cell types, tissues, and sizes (see Methods). We accounted for some significant challenges faced in real-world data and evaluated the generalizability of the algorithm. In this work, the tested datasets span diverse applications - including but not limited to immunotherapy (17), spatial transcriptomics (18), cell type profiling (19), cell differentiation (10). For each dataset, we demonstrate how top *InGene* genes selected via structure-preserving sampling reveal relevant genes according to the cell type, disease association, and tissue of origin.

After applying the structure-preserving sampling followed by the binary classification pipeline (see Methods), we use the top 500 ranked *InGene* genes for further analysis. To maintain parity with the other feature ranking methods used for comparison purposes, we also utilize the top 500 ranked genes from those methods.

We evaluated the 500 leading genes ranked by *InGene*, CV^2^, Fano Factor, Gini Index, PCA, MAST, and scGeneFit for the single-cell melanoma dataset. We found that the disease-gene association for *InGene* genes was most accurate compared to the other techniques (Figure 5).

The disease-gene association was captured from DisGeNET database (20) using EnrichR (21). *InGene* captures key genes associated with melanoma such as - ASAH1, GNG10, HSPG2, TMSB10, XAGE1A, DSG2, etc.

- ASAH1 is necessary for melanoma tumour growth and metastasis (22; 23) It was recently established in a study (22) that ASAH1 is overexpressed in melanoma and acts as a metabolic driver. Phenotypic plasticity, the ability of cells to change their characteristics, is an essential factor in developing drug resistance and metastasis in melanoma. The plasticity involves the regulation of different transcriptional programs, including MITF, and allows melanoma cells to switch between proliferative and invasive states. The protein ASAH1, which controls the metabolism of sphingolipids, was found to play a role in this switch in melanoma cells (22). Experiments showed that the ASAH1 could be used as a target for melanoma therapy, as it increases the effectiveness of BRAF inhibitors (22).
- GNG10, present in the family of G-proteins, is mutated in melanoma (24). According to a study published in 2010, GNG10 was found to have a high rate of non-synonymous mutations (changes to the DNA sequence that alter the amino acid sequence of the protein) in melanoma tumours. Moreover, these mutations were associated with the progression of the disease (24). The finding suggests that GNG10 may be involved in the development and progression of melanoma and that these mutations may be useful as markers for predicting the severity of the disease.
- HSPG2 is frequently mutated in melanoma and is over-expressed (25). Immune checkpoint inhibitor (ICIs) therapy improves the survival outcome of advanced melanoma patients. Zhang et. al (26) showed an association between melanoma patients with HSPG2 mutations had an improved ICI outcome. The authors found melanoma patients with mutations in the HSPG2 gene had better outcomes when treated with immunotherapy (ICI) compared to other patients. They also found that these patients had higher levels of immune cells that respond to cancer (response immunocytes), lower levels of immune cells that suppress the immune response (suppression immunocytes), more mutations in their cancer cells, and increased activity in pathways related to the immune system’s response to infection (interferon response-relevant signaling pathways). Based on these findings, the authors concluded that HSPG2 mutations may be a predictor of a good response to immunotherapy and may be used to guide treatment decisions for melanoma patients.
- TMSB10 is a well-established gene signature for predicting survival of metastatic melanoma patients (27; 28). The authors of a study (29) found that the protein TMSB10 was associated with the ability of melanoma cells to spread to other parts of the body (metastasis) in animal models and in fresh samples of human melanoma tissue. They also found that TMSB10 was not consistently related to the spread of cancer or the severity of disease in other types of cancer. Based on these findings, the authors concluded that TMSB10 might be a helpful marker for predicting the severity of cutaneous melanoma.
- DSG2 promotes vasculogenic mimicry (VM) in melanoma (30). Overexpression of DSG2 is associated with poor clinical outcomes in melanoma patients (31; 30). The study (30) establishes that DSG2 is expressed in primary and metastatic melanoma tissue but not in normal melanocytes. DSG2 plays a critical role in regulating the angiogenic activity of Endothelial Cells (ECs) and circulating endothelial progenitor cells (EPCs). The authors (30) have shown that DSG2 plays a similar cell-intrinsic role in melanoma by regulating VM and thus may prove to be a new treatment approach for melanoma.

### 2.3. *InGene* captures spatially differential genes

The spatial organization and heterogeneity of gene expression within a tissue have significant biological effects on the properties of the tissue. Regular transcriptome analyses using bulk or single-cell sequencing do not capture high-resolution spatial heterogeneity. Using these techniques, the rich spatial information about gene expression is lost. Recent developments in spatial transcriptomics capture spatial information by using DNA barcodes to distinguish different spots in the tissue. Detecting spatially differentially expressed genes can help reveal new marker genes, pathways, and molecular mechanisms and could also prove to be therapeutic targets. We used the spatial expression profile of the publicly available spatial gene expression for Human Breast Cancer (Block A Section 1) from the 10X genomics support website (18) Notably, *InGene* has the highest overlap with differentially expressed genes from SpatialDE (32), a well-known method specially designed to reveal differentially expressed genes in spatial transcriptomics data (Figure S12[E]).

*InGene* successfully captured relevant differentially expressed genes in an unsupervised manner without any input of spatial information. Some relevant differential genes captured by *InGene* are - MYC, EIF4A2, HSPG2, JPT1, NSMCE2.

- MYC is a protein that plays a role in controlling the growth and division of cells. When MYC is not regulated correctly, it can contribute to the development of various types of cancer, including breast cancer. MYC has been found to be active in the progression of breast cancer, and when combined with the loss of another protein called BRCA1, it can lead to the development of a specific type of breast cancer called basal-like breast cancer. MYC may also contribute to resistance to treatment for cancer, as it is influenced by the estrogen receptor and epidermal growth factor receptor pathways (33; 34).
- EIF4A2 plays a role in mRNA translation and has been implicated in cancer development and progression of breast cancer (35). According to a study by Liu et al., eIF4A2 mRNA levels were significantly higher in breast cancer tissues resistant to paclitaxel treatment compared to those that were sensitive to paclitaxel (36). Additionally, experiments have shown that reducing the amount of eIF4A2 protein in triple-negative breast cancer cells led to a decrease in cell proliferation and an increase in cell death (apoptosis) (36). These findings suggest that eIF4A2 may be a potential target for breast cancer treatment, particularly for triple-negative breast cancer.
- HSPG2, also known as perlecan, is a protein involved in the extracellular matrix and overexpressed in some types of cancer, including breast cancer (37; 38). Several studies have demonstrated that HSPG2 overexpression was associated with invasion, metastasis, and an inferior survival outcome in triple-negative breast cancer (39). HSPG2 has been reported as a novel target for breast cancer (40; 41).
- JPT1, also known as HN1 was found to be upregulated in breast cancer tissues (42). The authors of the study concluded that HN1 may contribute to the progression of breast cancer by increasing the expression of the protein MYC and that it may be a potential target for the treatment of breast cancer. Another study published in 2018 found that the HN1-like gene (HN1L) is frequently altered in breast cancer, particularly in triple-negative breast cancer (TNBC), and is associated with a poorer prognosis for patients (43). This study found that reducing the amount of HN1L protein in breast cancer cells led to a decrease in the population of breast cancer stem cells, made chemotherapy more effective, and slowed the progression of TNBC in animal models. Additionally, the researchers found that certain gene patterns were linked to shorter disease-free survival for TNBC patients. These findings suggest that HN1L may be a potential target for the treatment of TNBC.
- NSMCE2 was identified as a novel super-enhancer-associated gene – NSMCE2 in a recent study (44). The authors found that high levels of NSMCE2 are associated with a poor prognosis for breast cancer patients. They also found that disrupting certain regulatory regions called super-enhancers can increase the production of NSMCE2 and that high levels of NSMCE2 are linked to a poor response to chemotherapy, particularly in patients with aggressive triple-negative and HER2-positive breast cancer. Furthermore, decreasing the amount of NSMCE2 in breast cancer cells made them more sensitive to chemotherapy treatment.

### 2.4. *InGene* as an alternative unsupervised method to obtain differential genes

The above results show that *InGene* efficiently reveals relevant genes by learning the factors that dominate the latent representation of the dataset. Therefore, *InGene* can be used as an alternative to popular unsupervised methods to obtain differentially expressed genes (DEGs), such as t-test and wilcoxon test. It should be noted that the popular methods for DEG selection for single-cell RNA-seq data are supervised in nature (MAST (15), scGeneFit (16)). Notably, existing DE gene selection methods have been restricted to two-group comparisons. This is a hindrance, especially when it comes to single-cell data, which often has multiple groups in the dataset. Comparing only two groups at a time leads to post-processing overhead. It also does not allow the user to look at the comprehensive overall result because taking a union of all DE genes from each two-group comparison is not an effective solution to this end. *InGene* overcomes the shortcoming by ranking DE genes representative of each group present in the dataset. We achieve so by converting the multi-class classification problem to a binary classification problem (see Methods). Thus, the user does not need to perform additional steps for a multi-group comparison when using *InGene*. We also demonstrate in our experiments that the top 500 *InGene* genes lead to a higher classification accuracy (adjusted rand index (ARI)). *InGene* works efficiently on both small and large datasets. We observed that *InGene* genes efficiently describe the associated cell types for both PBMC68K (68000 cells) and human brain data (466 cells) (19; 10).

## 3. Methods

### 3.1. Datasets

To evaluate *InGene*, we applied the method to four publicly available single-cell datasets spanning different sizes, cell types, and diseases. The following section provides a brief description.

We used single-cell RNA sequencing (scRNA-seq) of Melanoma patient tumours (17) comprising 4,645 single cells, profiling malignant, immune, stromal and endothelial cells. Post filtering, 11217 genes and 4513 were retained. We also tested if *InGene* is capable of capturing spatially differential genes. To this regard, spatial gene expression of breast cancer specimens profiled via STseq using the Visium platform of 10x Genomics (Pleasanton, CA, USA) (18) consisting of 3813 spots, comprising basal, stroma, macrophage, luminal, mesenchymal, endothelial. B-Cell and T-cells were used. Post filtering 3209 genes and all the spots were retained.

We used two additional datasets to determine the scalability of *InGene*. The first is scRNA-seq data consisting of approximately 68000 fresh peripheral blood mononuclear cells (PBMCs) collected from a healthy donor (19). Single-cell expression profiles of 11 purified sub-populations of PBMCs are used as a reference for cell type annotation. Next, we wanted to determine if the algorithm is effective on datasets with a few cells. Darmanis et al. used single-cell RNA sequencing on 466 cells to capture human brain transcriptome diversity at the single-cell level (10). We filtered the three most prevalent cell types, namely neurons, astrocytes and oligodendrocytes, for the assessment. We were left with 227 cells and 14692 genes post-filtering.

### 3.2. Preprocessing of scRNA-seq data

Given a raw count data matrix, poorly expressed genes were first filtered out. Cells with a low number of detected genes were also ignored. A cell needs to have at least 200 genes expressed to be retained. Similarly, genes that were not expressed in more than one percent of the cells were filtered out. The thresholds may be modified depending on the size and dimension of the dataset. Seurat was used for data normalization and scaling. Of note, we did not limit scaling the data with respect to only the top variable genes (the default is top 2000 CV^2^ genes in Seurat). Instead, we chose to retain all the genes post-filtering.

### 3.3. Gaussian mixture modeling of single cells and structure preserving sampling

Once the count matrix is filtered, normalized and scaled, we utilize the Seurat pipeline to cluster the dataset using the Leiden algorithm (45). The clustered data is then projected on the latent dimensions (tSNE/UMAP). Once we obtain the two-dimensional map, we hypothesize that each single cell x is populated from a mixture of k bivariate Gaussians,

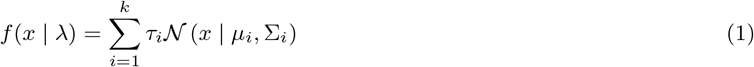

where *τ*_*i*_, *i* = 1, …, *k*, are mixture weights that add up to 1, *N* (*x* | *μ*_*i*_, Σ_*i*_), *i* = 1, …, *k*, are the components of bivariate Gaussian densities, each of which is characterized by a mean vector *μ*_*i*_ and covariance matrix Σ_*i*_.

These parameters are collectively represented by *λ*, where *λ* = *{τ*_*i*_, *μ*_*i*_, Σ_*i*_*}* . The value of *k* is equivalent to the number of clusters determined by the Leiden algorithm. The values of *μ*_*i*_ and Σ_*i*_ are calculated in a supervised manner for each cluster *i* using the MclustDA function from the mclust package (46). Thus, with a given *μ*_*i*_ and Σ_*i*_ for a cluster *i*, we assign each cell of the cluster *i* a probability or likelihood of belonging to it. We then subsample cells from each cluster by leveraging the likelihood values. If a cluster *i* contains *n* cells, we choose *min*(*p, n*) cells with the highest likelihood as well as *min*(*p, n*) cells with the lowest likelihood of belonging to cluster *i*, where the value of *p* typically ranges between 10 to 20. In parallel, we perform Support Vector Machine (SVM) (47) classification using the one-against-all (one-vs-rest) strategy. Given *k* clusters, *k* binary classifiers are generated. Each classifier is responsible for distinguishing a cluster *i* from the remaining clusters. Let us consider that cells belonging to the current cluster *i* are given the label +1, and cells belonging to all the clusters are given the label *−*1. SVM looks for a hyperplane (Equation 2), which separates the data from classes *y*_**i**_ (+1 and *−*1) with a maximal margin.

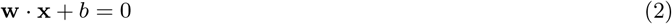

where **w** is the normal vector to the hyperplane and *b* is an offset. The margin maximization solves a convex quadratic optimization problem, in which the resulting decision boundary or hyperplane is given by Equation 3:

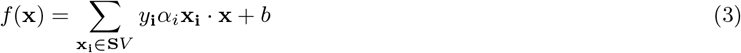

where the constants *α*_*i*_ are called Lagrange multipliers and are determined in the optimization process. Here SV corresponds to the set of support vectors, samples for which the associated Lagrange multipliers are larger than zero. These samples are those closest to the optimal hyperplane are the ones we choose to focus on. We select *min*(*p/*2, *s*) SVs from cluster *i* with the highest likelihood, where *s* is the total number of support vectors belonging to cluster *i*. In this manner, we select a few representative cells for each cluster.

Such a selection approach enables us to implement a structure-preserving subsampling. The method also allows for the efficient integration of cells across all clusters. The idea is that the cells that are the support vectors will represent the boundary or marginal cells, and the cells with the highest likelihood shall represent centrally located cells. Thus, features learnt from these cells together should be representative of the whole cluster. Since we are performing subsampling with only a few cells per cluster (approximately 20 to 30), the method is size agnostic and can scale well. Once we have all the selected cells, we construct a binary classification problem to score each gene.

### 3.4. Construction of a supervised binary classification problem

A new strategy was used to identify the key genes from a low-dimensional projection of scRNA-seq data. Instead of contrasting each group of cells to the rest, we modelled the differences between pairs of nearby cells to pairs of cells that were far off, regardless of their clusters of origin. A supervised bi-class classification problem was constructed for the same.

From each cluster, *i*, we utilize the subsampled cells from it. Input space for the classification problem consisted of a total of 2*M* data points, where each data point *d*_*i*_ *∈* ℝ ^|*G*|^ was created from a pair of cells (*m, n*). *G* here denotes the set of all expressed genes. Values of *d*_*i*_ are determined as follows.

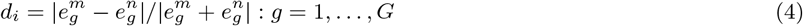

where 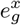 and 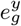 are expression of a gene *g ∈ G* in cells *m* and *n* respectively. Each data point *d*_*i*_ was assigned a label *y*_*i*_ *∈ {N, F }*, following a simple scheme. If both cells in a pair associated with a certain data point *d*_*i*_ shared the same cluster identity, *y*_*i*_ was set *N*, whereas *F* otherwise. We constructed *M* data points of each type to balance the data.

The above-described transformation is a unique way to link high dimensional space spanned by several thousands of genes to a corresponding latent space. We hypothesized that genes that best explain the proximity among pairs of cells in a latent space are the ones that influence the process of dimension reduction the most.

### 3.5. Gene selection using Random Forest

For classification, we chose the RandomForests (RF) algorithm (48) for the following reasons: 1. RF provides a Gini importance score for each variable (a gene in this case) by observing its contribution to constructing several decision trees on bootstrap training sets. 2. Any parametric assumptions do not substantiate it. 3. RF is a non-linear classifier. 4. The transformation depicted by Equation 4 results in folded distributions which may turn out to be incompatible with regression-based methods. For example, suppose the log-transformed expression of a gene follows a normal distribution. In that case, the absolute difference in its expression between a pair of cells will follow a folded normal distribution. Regression-based methods, in this scenario, are not guaranteed to produce stable results. To build the RF models, we employed a fast and memory-efficient implementation of the algorithm, distributed through an R package called ranger (49). Out Of Bag Error (OOBE) reported by ranger was considered as a measure of the goodness of a model. In our cases, OOBE varied between 2-5%. Finally, all expressed genes were prioritized based on the Gini importance scores. Important to note that RF models were never used for class prediction. Our main objective was to obtain the importance scores for a supplied set of genes, which was attained with the completion of the training phase.

### 3.6. Evaluation of genes

Suppose top genes suggested by *InGene* are principal determinants of the low-dimensional map of single-cell transcriptomes. In that case, we hypothesize that relative Euclidean distances between data points spanned by these genes would be preserved in the low-dimensional space. Spearman’s rank correlation coefficient between Euclidean distances was computed across cell pairs. Given a set of influential candidate genes, this simple measure comprehends their ability to reconstruct the distances observed in a latent space. For ease of interpretation, we termed this measure Reconstruction Accuracy or RA. We compared four additional unsupervised gene ranking methods (namely - CV^2^, PCA, Fano Factor, Gini Index) and two supervised (MAST and scGeneFit (15; 16)) gene ranking methods with *InGene* in this regard. We extract the top 1000 genes as per the rankings determined by each of these methods and compare their RA against the top 1000 *InGene* genes. RA scores were also computed on randomly sampled gene sets of length varying from 10 to 1000 over 20 iterations. This gives us a measurement of how the methods perform against random selection.

### 3.7. Assessing the relevance of selected genes

We selected 500 leading genes from each of the following methods: CV^2^, Fano Factor, Gini Index, PCA, MAST, scGeneFit, and *InGene*. To evaluate the biological significance of these gene sets, we utilized techniques such as disease gene association, cell-type identification, and survival analysis. To identify the disease gene association and cell types, we utilized the EnrichR web server. EnrichR is a widely used tool that allows users to input a list of genes and then extracts information from various well-known databases to provide insights on the collective functions of the gene lists. For survival analysis, we employed another reputable web server, GEPIA 2. GEPIA 2 can perform survival analysis on multiple genes using both tumour and normal samples from the TCGA and GTEx databases. To examine if the 500 top-ranked genes from the GSE72056 (single-cell melanoma dataset) are indicative of survival analysis, we selected the SKCM database in GEPIA 2. We used the gene-set from each of the unsupervised methods CV^2^, Fano Factor, Gini Index, PCA, and *InGene* in GEPIA 2. The output from GEPIA 2 includes Kaplan Meier (KM) survival curves for each set of input sets. Also, it provides the combined significance (*p* value) of the gene sets in the survival of cancer patients for the chosen cancer type. For both EnrichR and GEPIA 2, we used the official gene symbols as input. Any gene symbols that did not align with the pre-existing symbols in the internal databases of EnrichR and GEPIA 2 were eliminated from the analysis. Figure 4 shows that *InGene* genes are predictive of survival for melanoma patients.

**Fig. 4:**
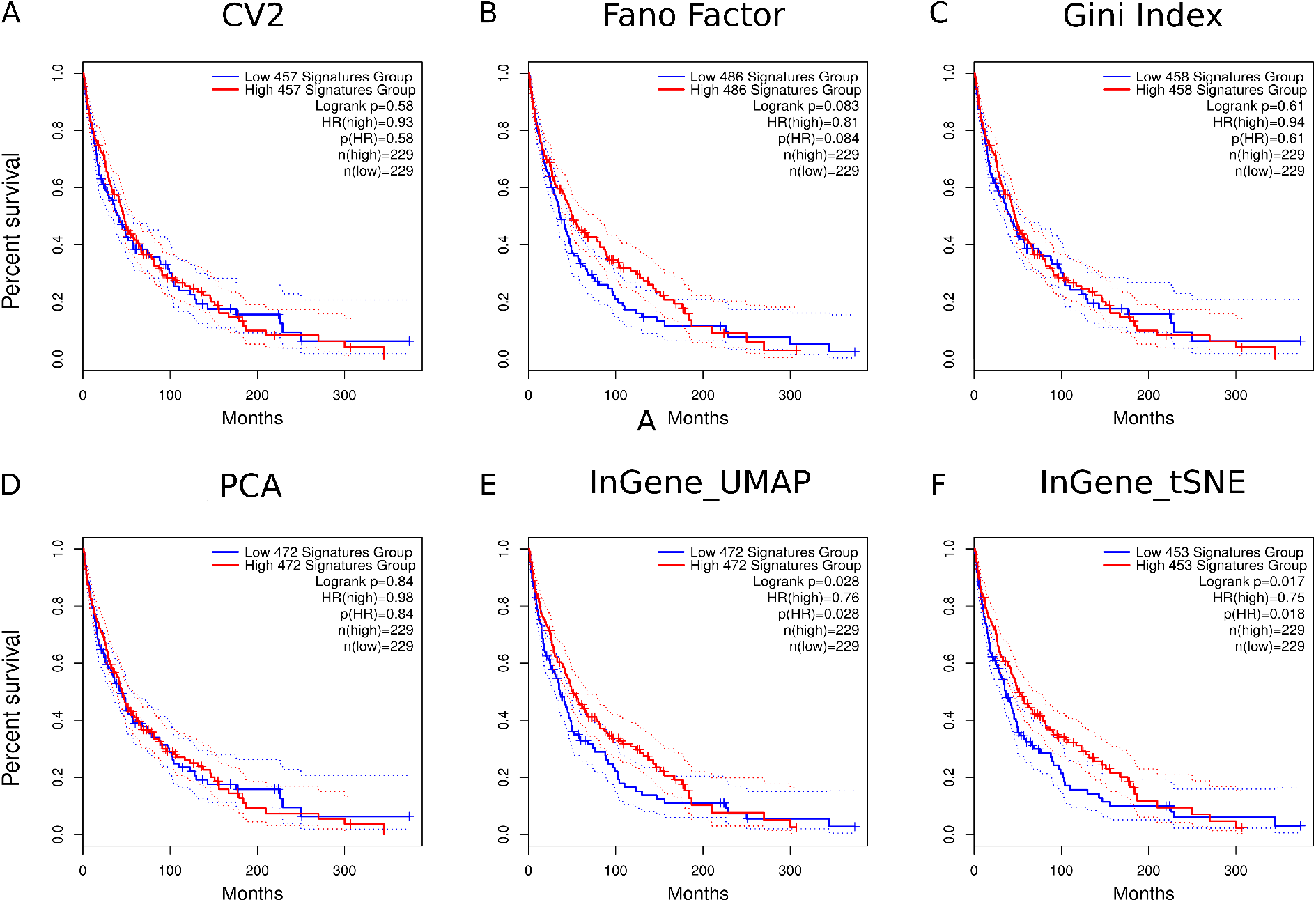
Kaplan-Meier (KM) plots for TCGA-SKCM dataset with top 500 genes obtained from each method. **(A)** Survival analysis (KM) on TCGA-SKCM dataset with top 500 CV2 genes obtained from melanoma dataset (GSE72056) *p*-value is 0.58. **(B)** Survival analysis (KM) on TCGA-SKCM dataset with top 500 Fano-Factor genes obtained from melanoma dataset (GSE72056) *p*-value is 0.08. **(C)** Survival analysis (KM) on TCGA-SKCM dataset with top 500 Gini genes obtained from melanoma dataset (GSE72056) *p*-value is 0.61. **(D)** Survival analysis (KM) on TCGA-SKCM dataset with top 500 PCA genes obtained from melanoma dataset (GSE72056) *p*-value is 0.84. **(E)** Survival analysis (KM) on TCGA-SKCM dataset with top 500 *InGene* genes obtained using 2D UMAP embeddings of melanoma dataset (GSE72056) *p*-value is 0.02. **(F)** Survival analysis (KM) on TCGA-SKCM dataset with top 500 *InGene* genes obtained using 2D t-SNE embeddings of melanoma dataset (GSE72056) *p*-value is 0.017.

**Fig. 5:**
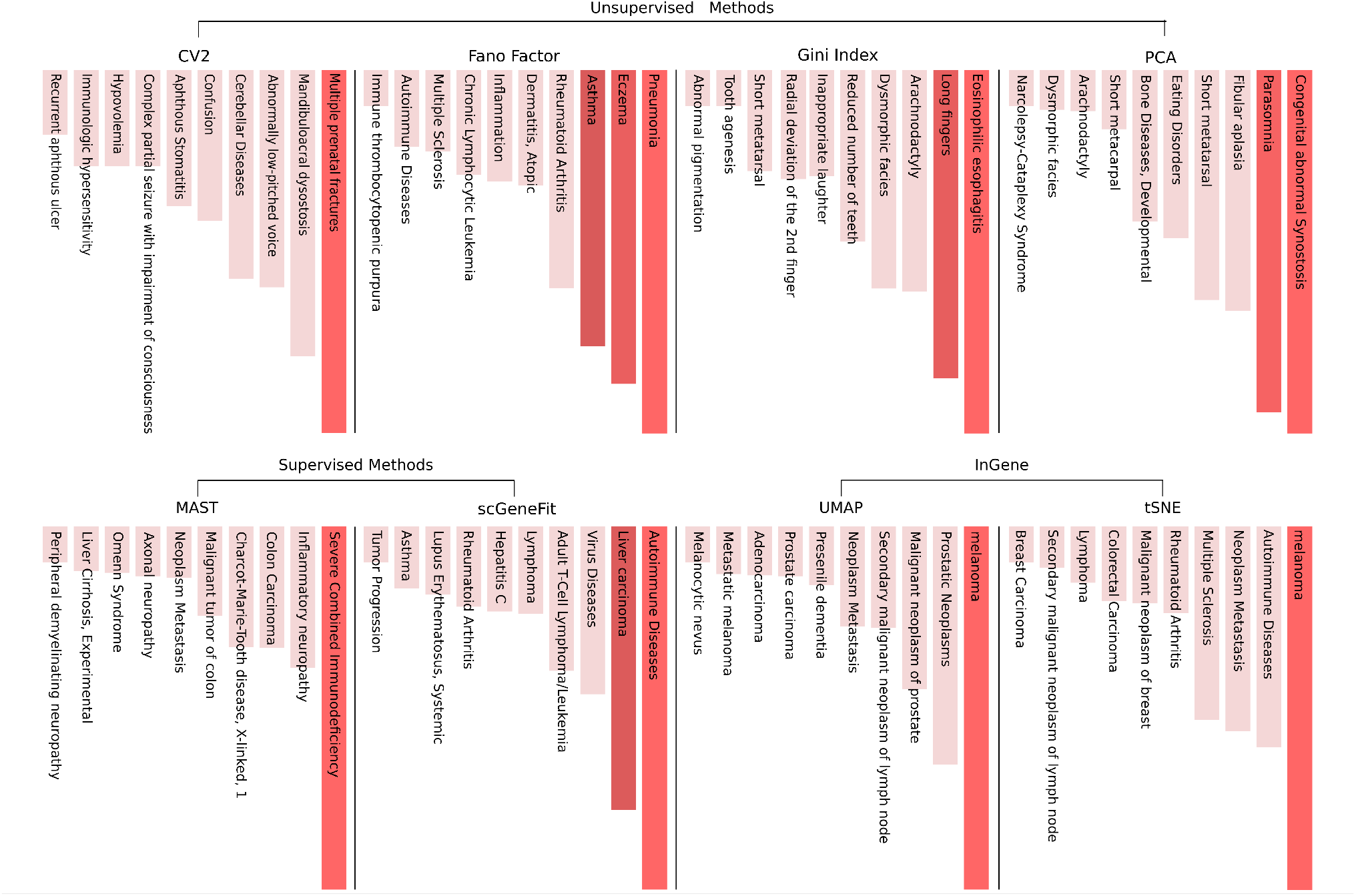
Disease-Gene association for top 500 genes from each method for GSE72056. *InGene* genes obtained from 2D UMAP and t-SNE embeddings correctly rank Melanoma as the associated disease, with the highest significance. Other methods - both supervised and unsupervised fail to capture the disease-gene association for the dataset.

**Fig. 6:**
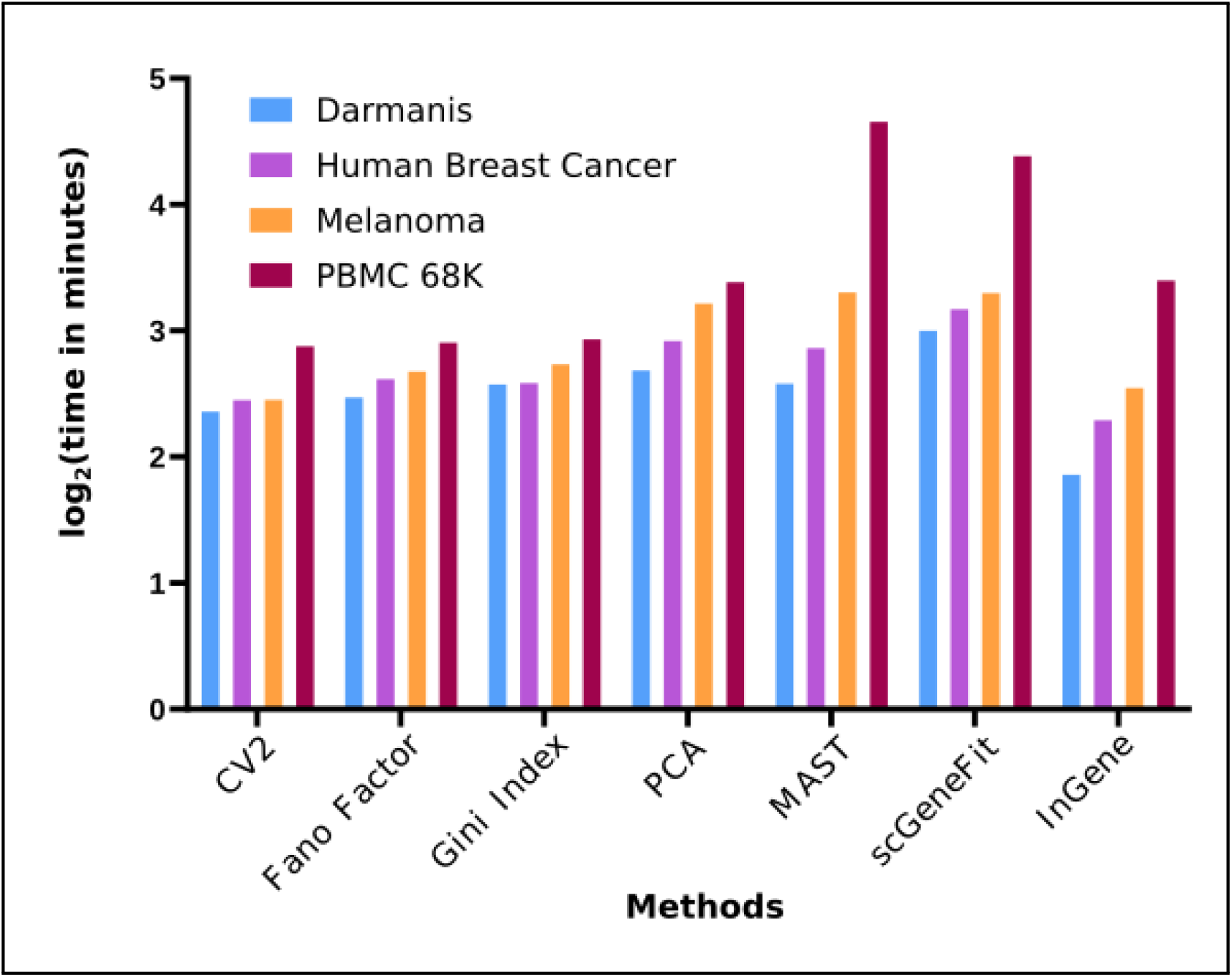
*InGene* is scalable. Run time recorded for each method (CV2, Fano Facor, Gini Index, PCA, MAST, scGeneFit, *InGene*) while varying the number of cells from *∼*200 to *∼*68K.

Noteworthy, we also assessed the speed and scalability of *InGene*. As the number of cells increase, the time taken by *InGene* increases linearly 6.

## 4. Discussion and Conclusion

*InGene* offers a robust method for analyzing scRNA-seq data without the need for pre-labeled training data and innovatively assigns importance scores to genes, enhancing understanding of cellular heterogeneity and disease-related gene identification. While these are crucial areas to address, the limitations of *InGene* must also be discussed. Computational methods like *InGene* may face challenges in accurately interpreting data due to the inherent biological variability among samples. This variability can arise from differences in tissue types, disease states, or even individual genetic differences. Since *InGene*’s accuracy is contingent upon the effectiveness of UMAP and tSNE, any limitations or inaccuracies inherent in these methods directly impact *InGene*’s performance. Considering that no dimensionality reduction technique is flawless, especially in the context of complex and often noisy biological data, this reliance can potentially be a limitation. Given these complexities, extensive validation across a wide range of datasets is crucial to ensure the robustness and generalizability of *InGene. InGene*, in principle, can be applied to any linear or nonlinear dimension reduction method to extract relevant genes. *InGene* can also be used as an alternative to differential expression analysis since :

- The associated gene ranking method is free of any parametric assumption, which makes *InGene* assay-independent.
- *InGene* does not require each group of cells to be compared against the rest. Instead, it poses the whole problem of cell type-specific gene finding as a single bi-class classification problem.

*InGene* tackles the variability and complexity of gene expression in cancer, offering a more nuanced understanding at a cellular level. We now progress to a patient-centric approach in the next chapter. We pave the way for developing novel diagnostic methods crucial for early and accurate cancer detection, thus encompassing a broader spectrum of cancer patient care.

## Author Contributions

D.S. conceived the project. C.G. designed the algorithm and the experiments. C.G. implemented the algorithm and performed all the experiments. N.B. contributed to preparing the manuscript and editing the draft. All the authors discussed the results, co-wrote and reviewed the manuscript.

## Acknowledgements

The authors thank IIIT Delhi for providing the resources to carry out this project.

## Availability and Implementation

*InGene* software package is available at: https://github.com/cgiiitd/InGene. The package can be installed in R using `devtools::install_github`(“cgiiitd/InGene/R/InGene”)

## Data Availability

The study uses various publicly available scRNA-seq datasets. The PBMC gene expression dataset is freely available for download from (50). The melanoma dataset can be accessed at the GEO under accession code GSE72056. The gene expression of breast cancer specimens profiled via STseq using the Visium platform of 10x Genomics is available in the 10X website for download. The single-cell human brain data can be obtained from GEO under the accession code GSE67835.

## Declarations

There are no conflicts of interest.

## References

1. A. Jindal, P. Gupta, D. Sengupta et al., “Discovery of rare cells from voluminous single cell expression data,” Nature communications, vol. 9, no. 1, pp. 1–9, 2018.

2. M. Tellez-Gabriel, B. Ory, F. Lamoureux, M.-F. Heymann, and D. Heymann, “Tumour heterogeneity: the key advantages of single-cell analysis,” International journal of molecular sciences, vol. 17, no. 12, p. 2142, 2016.

3. R. Bacher and C. Kendziorski, “Design and computational analysis of single-cell rna-sequencing experiments,” Genome biology, vol. 17, no. 1, pp. 1–14, 2016.

4. M. Sewell, “Principal component analysis,” 2007.

5. L. Van der Maaten and G. Hinton, “Visualizing data using t-sne.” Journal of machine learning research, vol. 9, no. 11, 2008.

6. L. McInnes, J. Healy, and J. Melville, “Umap: Uniform manifold approximation and projection for dimension reduction,” arXiv preprint arXiv:1802.03426, 2018.

7. B. Treutlein, D. G. Brownfield, A. R. Wu, N. F. Neff, G. L. Mantalas, F. H. Espinoza, T. J. Desai, M. A. Krasnow, and S. R. Quake, “Reconstructing lineage hierarchies of the distal lung epithelium using single-cell rna-seq,” Nature, vol. 509, no. 7500, pp. 371–375, 2014.

8. Y. Wu, P. Tamayo, and K. Zhang, “Visualizing and interpreting single-cell gene expression datasets with similarity weighted nonnegative embedding,” Cell systems, vol. 7, no. 6, pp. 656–666, 2018.

9. E. Becht, L. McInnes, J. Healy, C.-A. Dutertre, I. W. Kwok, L. G. Ng, F. Ginhoux, and E. W. Newell, “Dimensionality reduction for visualizing single-cell data using umap,” Nature biotechnology, vol. 37, no. 1, pp. 38–44, 2019.

10. S. Darmanis, S. A. Sloan, Y. Zhang, M. Enge, C. Caneda, L. M. Shuer, M. G. Hayden Gephart, B. A. Barres, and S. R. Quake, “A survey of human brain transcriptome diversity at the single cell level,” Proceedings of the National Academy of Sciences, vol. 112, no. 23, pp. 7285–7290, 2015.

11. B. S. Everitt and A. Skrondal, “The cambridge dictionary of statistics,” 2010.

12. U. Fano, “Ionization yield of radiations. ii. the fluctuations of the number of ions,” Physical Review, vol. 72, no. 1, p. 26, 1947.

13. M. Wright Muelas, F. Mughal, S. O’Hagan, P. J. Day, and D. B. Kell, “The role and robustness of the gini coefficient as an unbiased tool for the selection of gini genes for normalising expression profiling data,” Scientific reports, vol. 9, no. 1, pp. 1–21, 2019.

14. S. Wold, K. Esbensen, and P. Geladi, “Principal component analysis,” Chemometrics and intelligent laboratory systems, vol. 2, no. 1-3, pp. 37–52, 1987.

15. G. Finak, A. McDavid, M. Yajima, J. Deng, V. Gersuk, A. K. Shalek, C. K. Slichter, H. W. Miller, M. J. McElrath, M. Prlic et al., “Mast: a flexible statistical framework for assessing transcriptional changes and characterizing heterogeneity in single-cell rna sequencing data,” Genome biology, vol. 16, no. 1, pp. 1–13, 2015.

16. B. Dumitrascu, S. Villar, D. G. Mixon, and B. E. Engelhardt, “Optimal marker gene selection for cell type discrimination in single cell analyses,” Nature communications, vol. 12, no. 1, pp. 1–8, 2021.

17. I. Tirosh, B. Izar, S. M. Prakadan, M. H. Wadsworth, D. Treacy, J. J. Trombetta, A. Rotem, C. Rodman, C. Lian, G. Murphy et al., “Dissecting the multicellular ecosystem of metastatic melanoma by single-cell rna-seq,” Science, vol. 352, no. 6282, pp. 189–196, 2016.

18. “10x genomics. human breast cancer (block a section 1),” 2019, https://support.10xgenomics.com/spatial-gene-expression/datasets/1.0.0/V1 Breast Cancer Block A Section 1.

19. “Fresh 68k pbmcs (donor a),” 2016, https://support.10xgenomics.com/single-cell-gene-expression/datasets/1.1.0/ fresh 68k pbmc donor a.

20. J. Piñero, À. Bravo, N. Queralt-Rosinach, A. Gutiérrez-Sacristán, J. Deu-Pons, E. Centeno, J. García-García, F. Sanz, and L. I. Furlong, “Disgenet: a comprehensive platform integrating information on human disease-associated genes and variants,” Nucleic acids research, p. gkw943, 2016.

21. E. Y. Chen, C. M. Tan, Y. Kou, Q. Duan, Z. Wang, G. V. Meirelles, N. R. Clark, and A. Ma’ayan, “Enrichr: interactive and collaborative html5 gene list enrichment analysis tool,” BMC bioinformatics, vol. 14, no. 1, pp. 1–14, 2013.

22. P. Malvi, R. Janostiak, A. Nagarajan, X. Zhang, and N. Wajapeyee, “N-acylsphingosine amidohydrolase 1 promotes melanoma growth and metastasis by suppressing peroxisome biogenesis-induced ros production,” Molecular metabolism, vol. 48, p. 101217, 2021.

23. J. Leclerc, D. Garandeau, C. Pandiani, C. Gaudel, K. Bille, N. Nottet, V. Garcia, P. Colosetti, S. Pagnotta, P. Bahadoran et al., “Lysosomal acid ceramidase asah1 controls the transition between invasive and proliferative phenotype in melanoma cells,” Oncogene, vol. 38, no. 8, pp. 1282–1295, 2019.

24. L. I. Cárdenas-Navia, P. Cruz, J. C. Lin, N. C. S. Program, S. A. Rosenberg, Y. Samuels et al., “Novel somatic mutations in heterotrimeric g proteins in melanoma,” Cancer biology & therapy, vol. 10, no. 1, pp. 33–37, 2010.

25. I. R. Cohen, A. D. Murdoch, M. F. Naso, D. Marchetti, D. Berd, and R. V. Iozzo, “Abnormal expression of perlecan proteoglycan in metastatic melanomas,” Cancer research, vol. 54, no. 22, pp. 5771–5774, 1994.

26. W. Zhang, Z. Lin, F. Shi, Q. Wang, Y. Kong, Y. Ren, J. Lyu, C. Sheng, Y. Li, H. Qin et al., “Hspg2 mutation association with immune checkpoint inhibitor outcome in melanoma and non-small cell lung cancer,” Cancers, vol. 14, no. 14, p. 3495, 2022.

27. C. Song, Z. Su, and J. Guo, “Thymosin β 10 is overexpressed and associated with unfavorable prognosis in hepatocellular carcinoma,” Bioscience reports, vol. 39, no. 3, 2019.

28. W. M. Hardesty, M. C. Kelley, D. Mi, R. L. Low, and R. M. Caprioli, “Protein signatures for survival and recurrence in metastatic melanoma,” Journal of proteomics, vol. 74, no. 7, pp. 1002–1014, 2011.

29. M. A. Weterman, G. N. Van Muijen, D. J. Ruiter, and H. P. Bloemers, “Thymosin β-10 expression in melanoma cell lines and melanocytic lesions: A new progression marker for human cutaneous melanoma,” International journal of cancer, vol. 53, no. 2, pp. 278–284, 1993.

30. L. Y. Tan, C. Mintoff, M. Z. Johan, B. W. Ebert, C. Fedele, Y. F. Zhang, P. Szeto, K. E. Sheppard, G. A. McArthur, E. Foster-Smith et al., “Desmoglein 2 promotes vasculogenic mimicry in melanoma and is associated with poor clinical outcome,” Oncotarget, vol. 7, no. 29, p. 46492, 2016.

31. W. K. Peitsch, Y. Doerflinger, R. Fischer-Colbrie, V. Huck, A. T. Bauer, J. Utikal, S. Goerdt, and S. W. Schneider, “Desmoglein 2 depletion leads to increased migration and upregulation of the chemoattractant secretoneurin in melanoma cells,” PLoS One, vol. 9, no. 2, p. e89491, 2014.

32. V. Svensson, S. A. Teichmann, and O. Stegle, “Spatialde: identification of spatially variable genes,” Nature methods, vol. 15, no. 5, pp. 343–346, 2018.

33. J. Xu, Y. Chen, and O. I. Olopade, “Myc and breast cancer,” Genes & cancer, vol. 1, no. 6, pp. 629–640, 2010.

34. Y. Fallah, J. Brundage, P. Allegakoen, and A. N. Shajahan-Haq, “Myc-driven pathways in breast cancer subtypes,” Biomolecules, vol. 7, no. 3, p. 53, 2017.

35. L. X. Yan, Q. N. Wu, Y. Zhang, Y. Y. Li, D. Z. Liao, J. H. Hou, J. Fu, M. S. Zeng, J. P. Yun, Q. L. Wu et al., “Knockdown of mir-21 in human breast cancer cell lines inhibits proliferation, in vitro migration and in vivotumor growth,” Breast cancer research, vol. 13, no. 1, pp. 1–14, 2011.

36. M. Liu, C. Gong, R. Xu, Y. Chen, and X. Wang, “Microrna-5195-3p enhances the chemosensitivity of triple-negative breast cancer to paclitaxel by downregulating eif4a2,” Cellular & Molecular Biology Letters, vol. 24, no. 1, pp. 1–11, 2019.

37. Z. Elgundi, M. Papanicolaou, G. Major, T. R. Cox, J. Melrose, J. M. Whitelock, and B. L. Farrugia, “Cancer metastasis: the role of the extracellular matrix and the heparan sulfate proteoglycan perlecan,” Frontiers in oncology, vol. 9, p. 1482, 2020.

38. R. V. Iozzo and R. D. Sanderson, “Proteoglycans in cancer biology, tumour microenvironment and angiogenesis,” Journal of cellular and molecular medicine, vol. 15, no. 5, pp. 1013–1031, 2011.

39. A. J. Giráldez, R. R. Copley, and S. M. Cohen, “Hspg modification by the secreted enzyme notum shapes the wingless morphogen gradient,” Developmental cell, vol. 2, no. 5, pp. 667–676, 2002.

40. S. Kalscheuer, V. Khanna, H. Kim, S. Li, D. Sachdev, A. DeCarlo, D. Yang, and J. Panyam, “Discovery of hspg2 (perlecan) as a therapeutic target in triple negative breast cancer,” Scientific reports, vol. 9, no. 1, pp. 1–11, 2019.

41. V. Khanna, S. Kalscheuer, J. Panyam, D. Yang, and S. Li, “Therapeutic efficacy of antibodies targeting domain 1 of hspg2 in breast cancer,” Cancer Research, vol. 78, no. 13 Supplement, pp. 2897–2897, 2018.

42. C. Zhang, B. Xu, S. Lu, Y. Zhao, and P. Liu, “Hn1 contributes to migration, invasion, and tumorigenesis of breast cancer by enhancing myc activity,” Molecular cancer, vol. 16, no. 1, pp. 1–10, 2017.

43. Y. Liu, D. S. Choi, J. Sheng, J. E. Ensor, D. H. Liang, C. Rodriguez-Aguayo, A. Polley, S. Benz, O. Elemento, A. Verma et al., “Hn1l promotes triple-negative breast cancer stem cells through lepr-stat3 pathway,” Stem cell reports, vol. 10, no. 1, pp. 212–227, 2018.

44. C. Di Benedetto, J. Oh, Z. Choudhery, W. Shi, G. Valdes, and P. Betancur, “Nsmce2, a novel super-enhancer regulated gene, is linked to poor prognosis and therapy resistance in breast cancer,” bioRxiv, 2022.

45. V. A. Traag, L. Waltman, and N. J. Van Eck, “From louvain to leiden: guaranteeing well-connected communities,” Scientific reports, vol. 9, no. 1, p. 5233, 2019.

46. L. Scrucca, M. Fop, T. B. Murphy, and A. E. Raftery, “mclust 5: clustering, classification and density estimation using gaussian finite mixture models,” The R journal, vol. 8, no. 1, p. 289, 2016.

47. M. A. Hearst, S. T. Dumais, E. Osuna, J. Platt, and B. Scholkopf, “Support vector machines,” IEEE Intelligent Systems and their applications, vol. 13, no. 4, pp. 18–28, 1998.

48. L. Breiman, “Random forests,” Springer Science and Business Media LLC, 2001. [Online]. Available: http://link.springer.com/10.1023/A:1010933404324

49. M. N. Wright and A. Ziegler, “ranger: A fast implementation of random forests for high dimensional data in c++ and r,” arXiv preprint arXiv:1508.04409, 2015.

50. “PBMC 68K datasetn,” https://github.com/10XGenomics/single-cell-3prime-paper/blob/master/pbmc68k_analysis/README.md.

