## Supplementary Information for "InGene: Finding influential genes from embeddings of nonlinear dimension reduction techniques"

, , 1–1
doi:

### InGene: Finding influential genes from embeddings of nonlinear dimension reduction techniques

**Chitrita Goswami[1] and Debarka Sengupta[1,2,3,4,*]**

[1]Department of Computer Science and Engineering, Indraprastha Institute of Information Technology, Okhla Industrial Estate, Phase III, 110020, Delhi, India, [2]Center for Computational Biology, Indraprastha Institute of Information Technology, Okhla Industrial Estate, Phase III, 110020, Delhi, India, [3]Center for Artificial Intelligence, Indraprastha Institute of Information Technology, Okhla Industrial Estate, Phase III, 110020, Delhi, India and [4]Institute of Health and Biomedicine Innovation, Queensland University of Technology, Australia

#### Abstract

#### Supplementary Figures

© The Author . Published

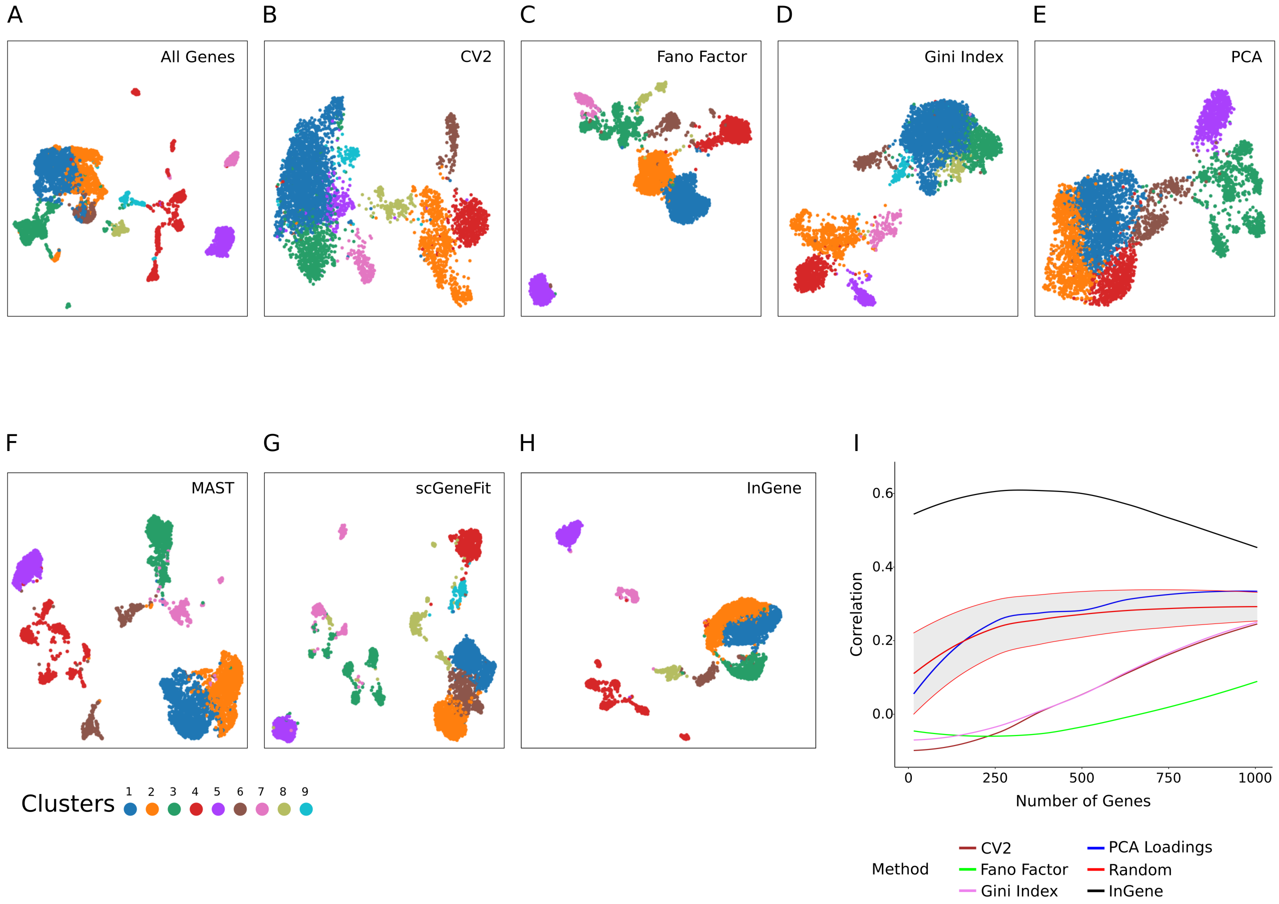


Fig. S 1: Explaining UMAP clusters for single-cell melanoma dataset with *InGene* **(A)** UMAP constructed with all the genes post-filtering. **(B)** UMAP constructed with top 500 CV2 genes. **(C)** UMAP constructed with top 500 Fano Factor genes. **(D)** UMAP constructed with top 500 Gini Index genes. **(E)** UMAP constructed with top 500 PCA genes. **(F)** UMAP constructed with top 500 MAST genes. **(G)** UMAP constructed with top 500 scGeneFit genes **(H)** UMAP constructed with top 500 *InGene* Factor genes. **(I)** Reconstruction Accuracy (RA) measured for leading 1000 genes for each method. Spearman's rank correlation measures the similarity between high-dimensional and low-dimensional distances. Correlation is calculated with the genes selected in high dimensional space for each method, and compared against randomly selected gene sets of length varying from 10 to 1000 over 20 iterations.

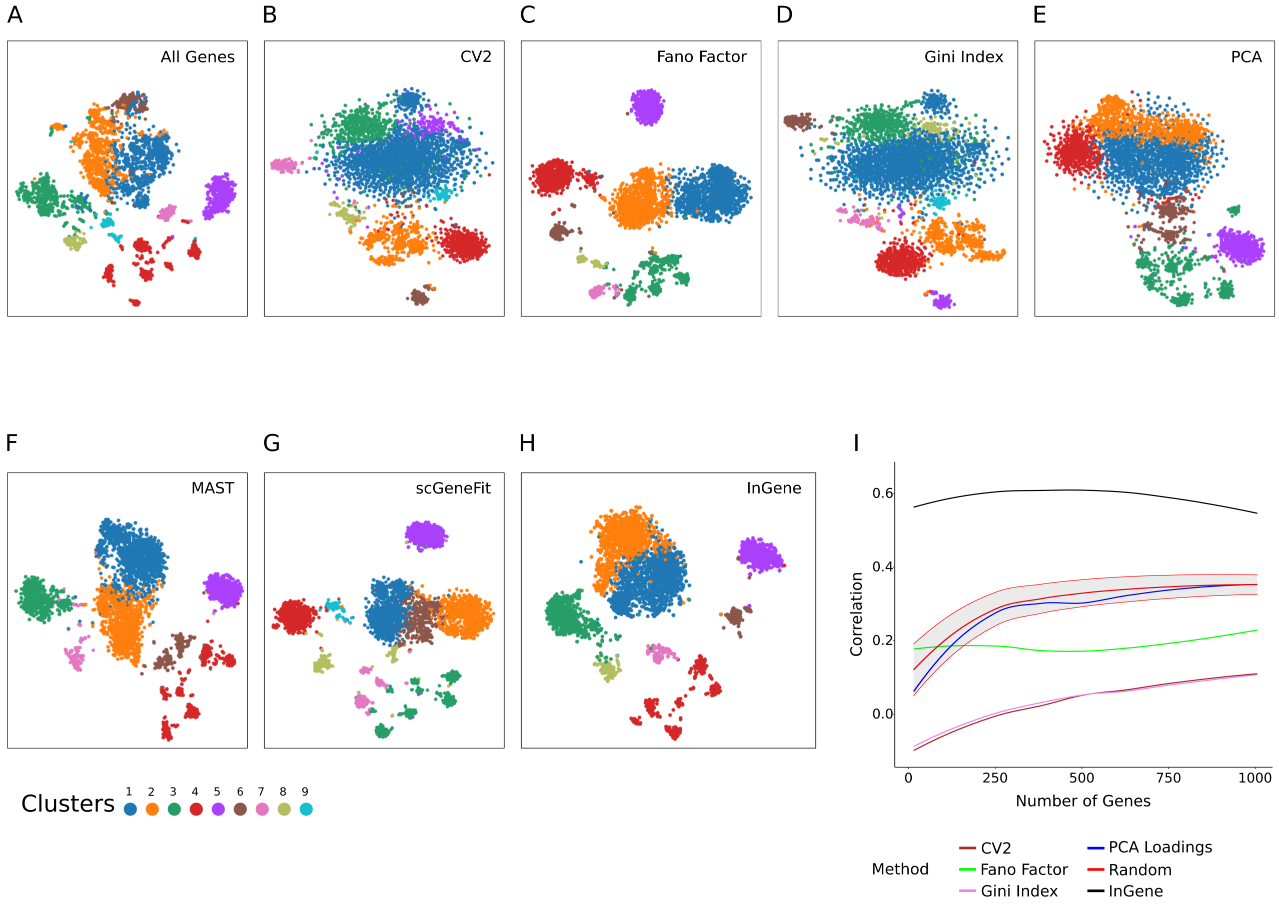


Fig. S 2: Explaining tSNE clusters for single-cell melanoma dataset with *InGene* **(A)** tSNE constructed with all the genes post-filtering. **(B)** tSNE constructed with top 500 CV2 genes. **(C)** tSNE constructed with top 500 Fano Factor genes. **(D)** tSNE constructed with top 500 Gini Index genes. **(E)** tSNE constructed with top 500 PCA genes. **(F)** tSNE constructed with top 500 MAST genes. **(G)** tSNE constructed with top 500 scGeneFit genes **(H)** tSNE constructed with top 500 *InGene* Factor genes. **(I)** Reconstruction Accuracy (RA) measured for leading 1000 genes for each method. Spearman's rank correlation measures the similarity between high-dimensional and low-dimensional distances. Correlation is calculated with the genes selected in high dimensional space for each method, and compared against randomly selected gene sets of length varying from 10 to 1000 over 20 iterations.

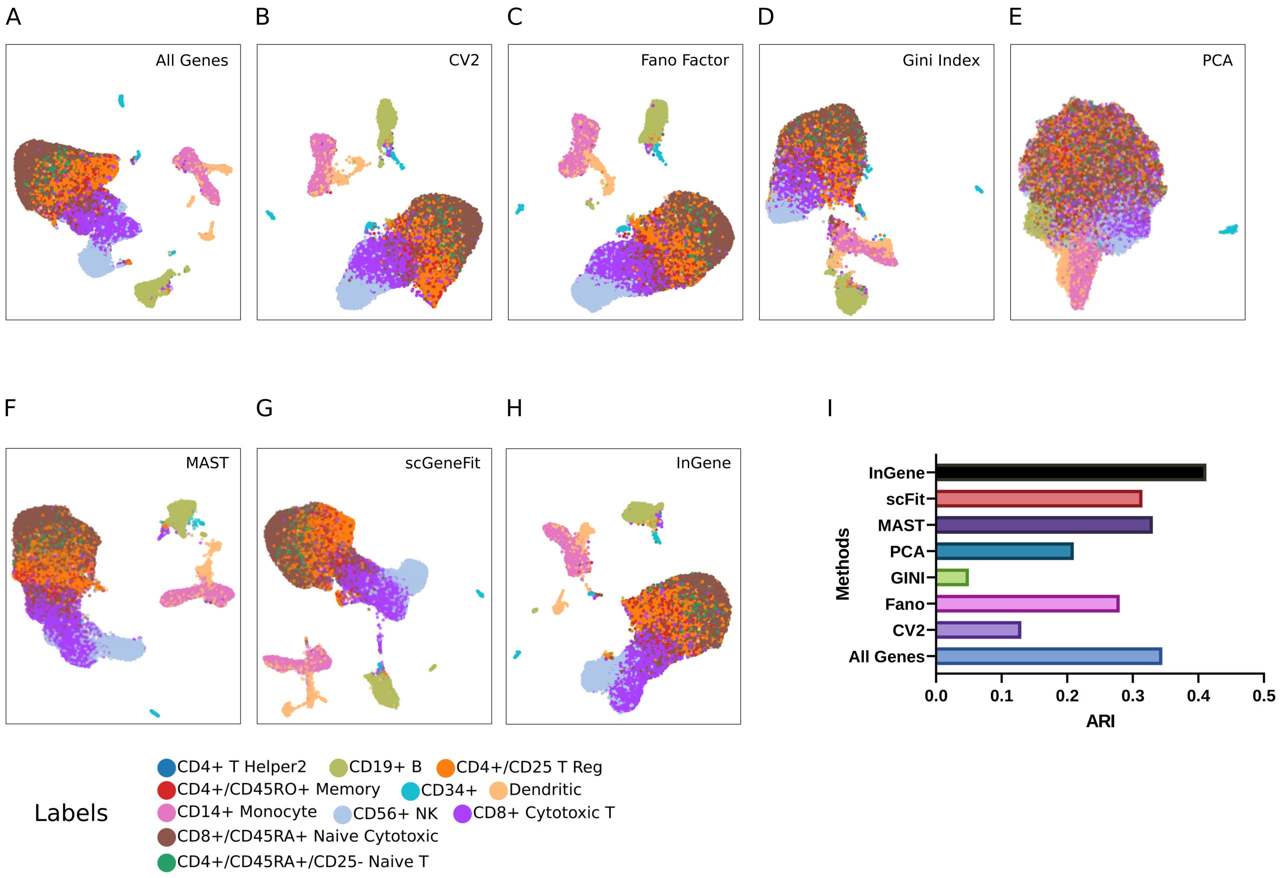


Fig. S 3: Explaining UMAP for single-cell PBMC 68K dataset with *InGene* **(A)** UMAP constructed with all the genes post-filtering. **(B)** UMAP constructed with top 500 CV2 genes. **(C)** UMAP constructed with top 500 Fano Factor genes. **(D)** UMAP constructed with top 500 Gini Index genes. **(E)** UMAP constructed with top 500 PCA genes. **(F)** UMAP constructed with top 500 MAST genes. **(G)** UMAP constructed with top 500 scGeneFit genes **(H)** UMAP constructed with top 500 *InGene* Factor genes. **(I)** ARI scores for the different methods. The gene set from each methods is used to cluster the dataset, using Leiden algorithm. The cluster labels obtained are then compared with the true labels to obtain the ARI values.

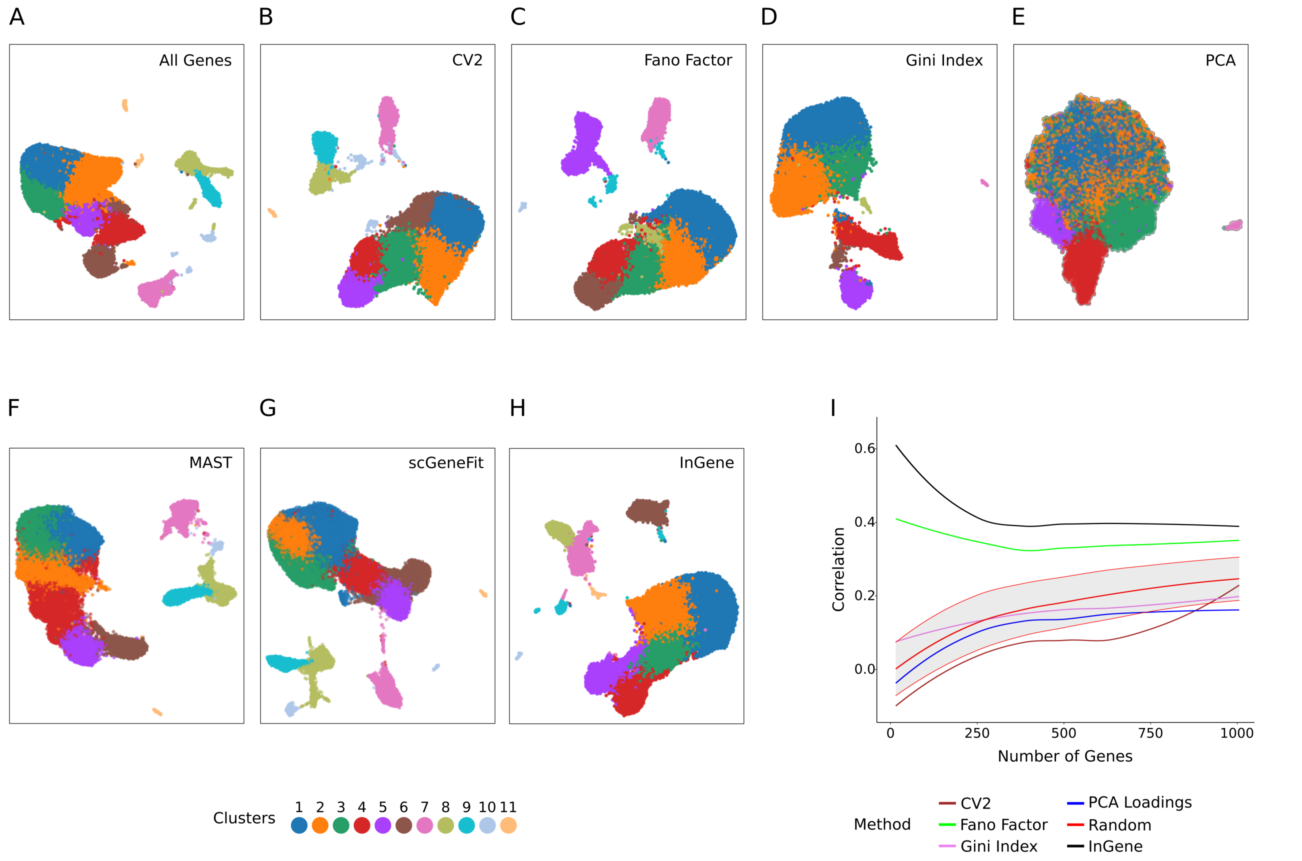


Fig. S 4: Explaining UMAP clusters for single-cell PBMC 68K dataset with *InGene* **(A)** UMAP constructed with all the genes post-filtering. **(B)** UMAP constructed with top 500 CV2 genes. **(C)** UMAP constructed with top 500 Fano Factor genes. **(D)** UMAP constructed with top 500 Gini Index genes. **(E)** UMAP constructed with top 500 PCA genes. **(F)** UMAP constructed with top 500 MAST genes. **(G)** UMAP constructed with top 500 scGeneFit genes **(H)** UMAP constructed with top 500 *InGene* Factor genes. **(I)** Reconstruction Accuracy (RA) measured for leading 1000 genes for each method. Spearman's rank correlation measures the similarity between high-dimensional and low-dimensional distances. Correlation is calculated with the genes selected in high dimensional space for each method, and compared against randomly selected gene sets of length varying from 10 to 1000 over 20 iterations.

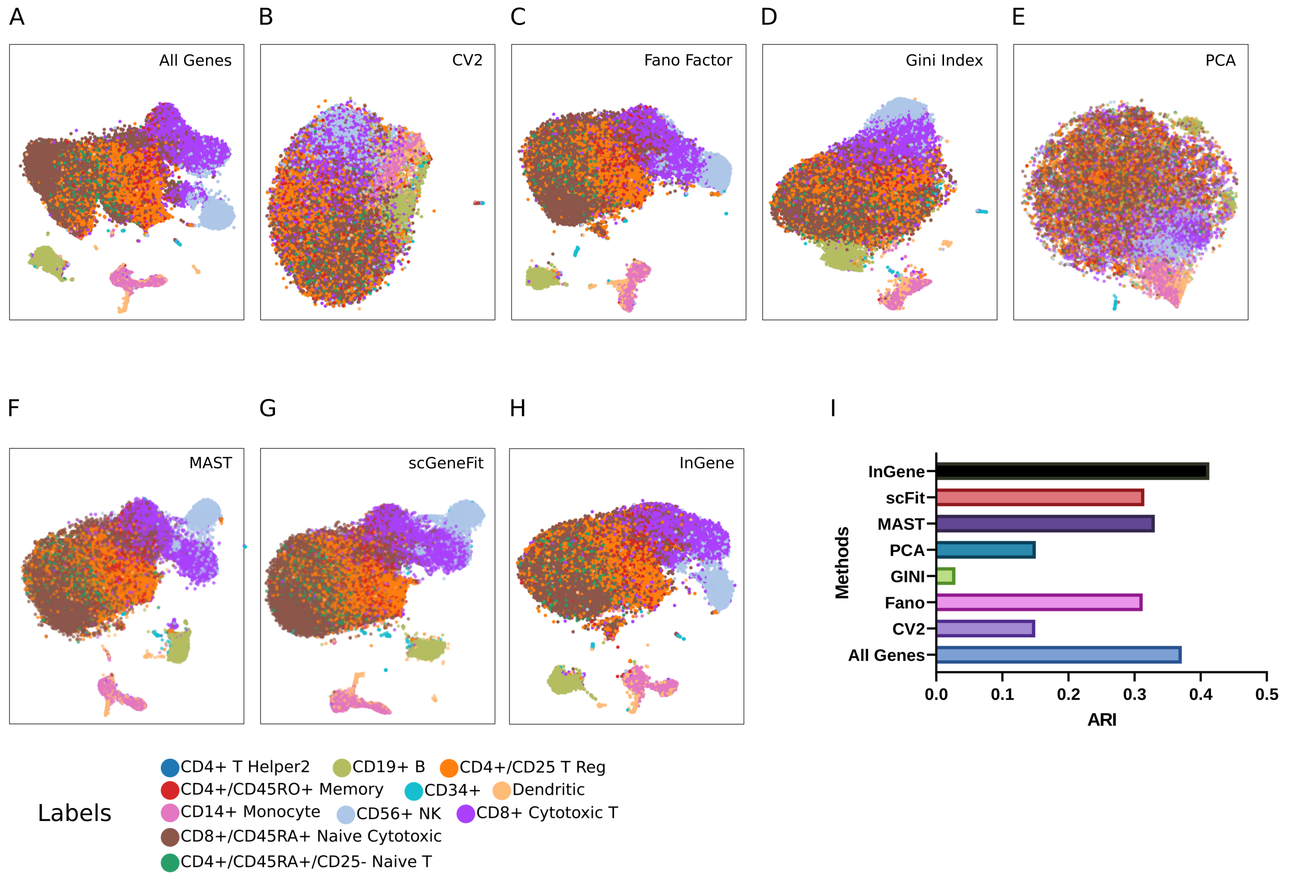


Fig. S 5: Explaining tSNE for single-cell PBMC 68K dataset with *InGene* **(A)** tSNE constructed with all the genes post-filtering. **(B)** tSNE constructed with top 500 CV2 genes. **(C)** tSNE constructed with top 500 Fano Factor genes. **(D)** tSNE constructed with top 500 Gini Index genes. **(E)** tSNE constructed with top 500 PCA genes. **(F)** tSNE constructed with top 500 MAST genes. **(G)** tSNE constructed with top 500 scGeneFit genes **(H)** tSNE constructed with top 500 *InGene* Factor genes. **(I)** ARI scores for the different methods. The gene set from each methods is used to cluster the dataset, using Leiden algorithm. The cluster labels obtained are then compared with the true labels to obtain the ARI values.

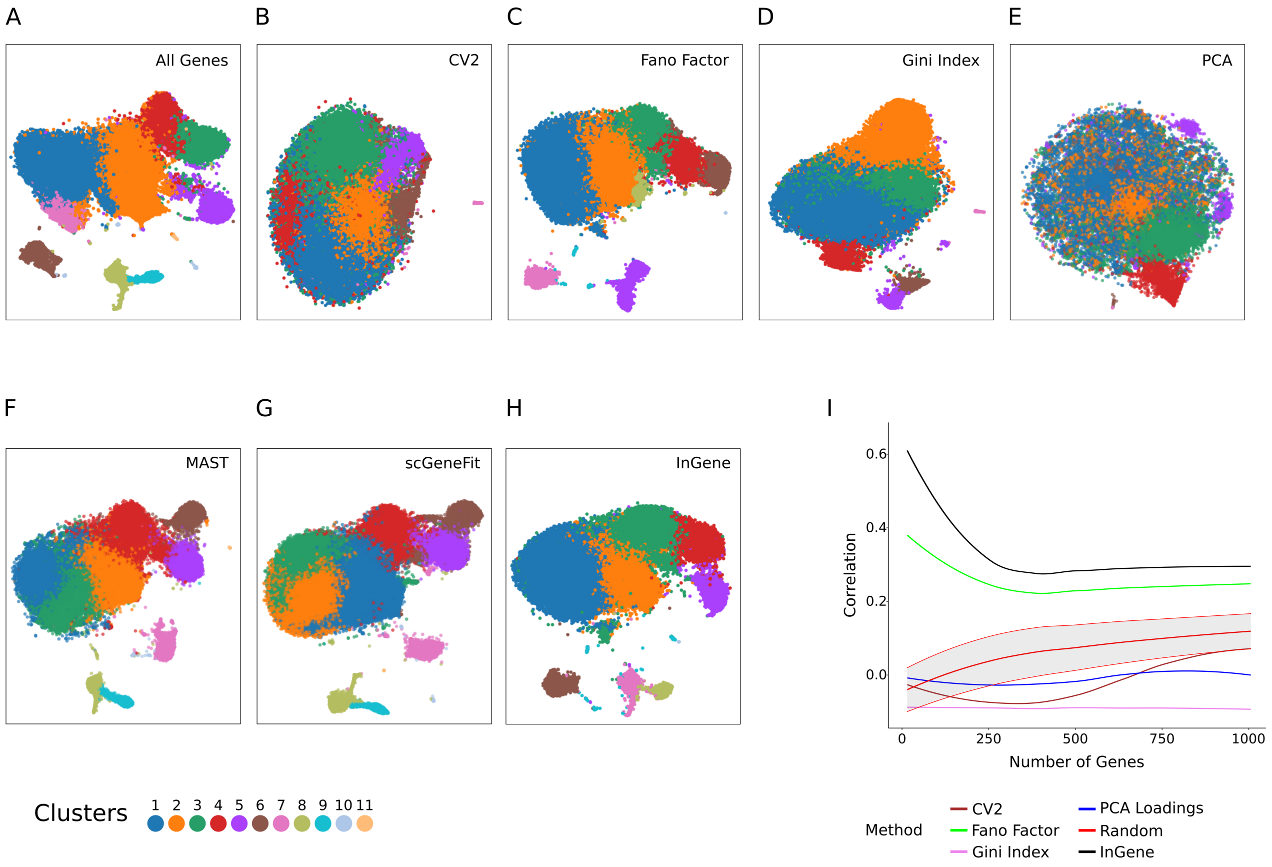


Fig. S 6: Explaining tSNE clusters for single-cell PBMC 68K dataset with *InGene* **(A)** tSNE constructed with all the genes post-filtering. **(B)** tSNE constructed with top 500 CV2 genes. **(C)** tSNE constructed with top 500 Fano Factor genes. **(D)** tSNE constructed with top 500 Gini Index genes. **(E)** tSNE constructed with top 500 PCA genes. **(F)** tSNE constructed with top 500 MAST genes. **(G)** tSNE constructed with top 500 scGeneFit genes **(H)** tSNE constructed with top 500 *InGene* Factor genes. **(I)** Reconstruction Accuracy (RA) measured for leading 1000 genes for each method. Spearman's rank correlation measures the similarity between high-dimensional and low-dimensional distances. Correlation is calculated with the genes selected in high dimensional space for each method, and compared against randomly selected gene sets of length varying from 10 to 1000 over 20 iterations.

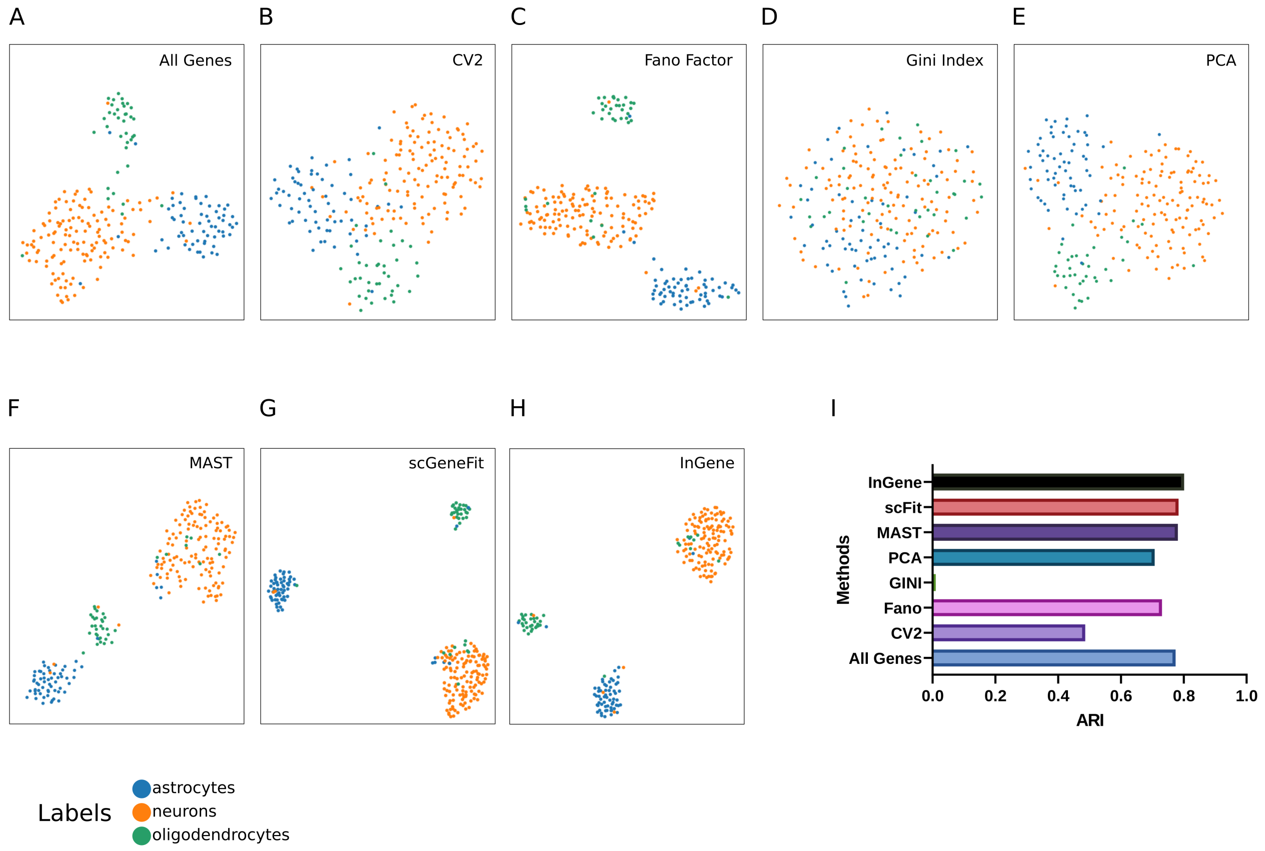


Fig. S 7: Explaining UMAP for single-cell Darmanis dataset with *InGene* **(A)** UMAP constructed with all the genes post-filtering. **(B)** UMAP constructed with top 500 CV2 genes. **(C)** UMAP constructed with top 500 Fano Factor genes. **(D)** UMAP constructed with top 500 Gini Index genes. **(E)** UMAP constructed with top 500 PCA genes. **(F)** UMAP constructed with top 500 MAST genes. **(G)** UMAP constructed with top 500 scGeneFit genes **(H)** UMAP constructed with top 500 *InGene* Factor genes. **(I)** ARI scores for the different methods. The gene set from each methods is used to cluster the dataset, using Leiden algorithm. The cluster labels obtained are then compared with the true labels to obtain the ARI values.

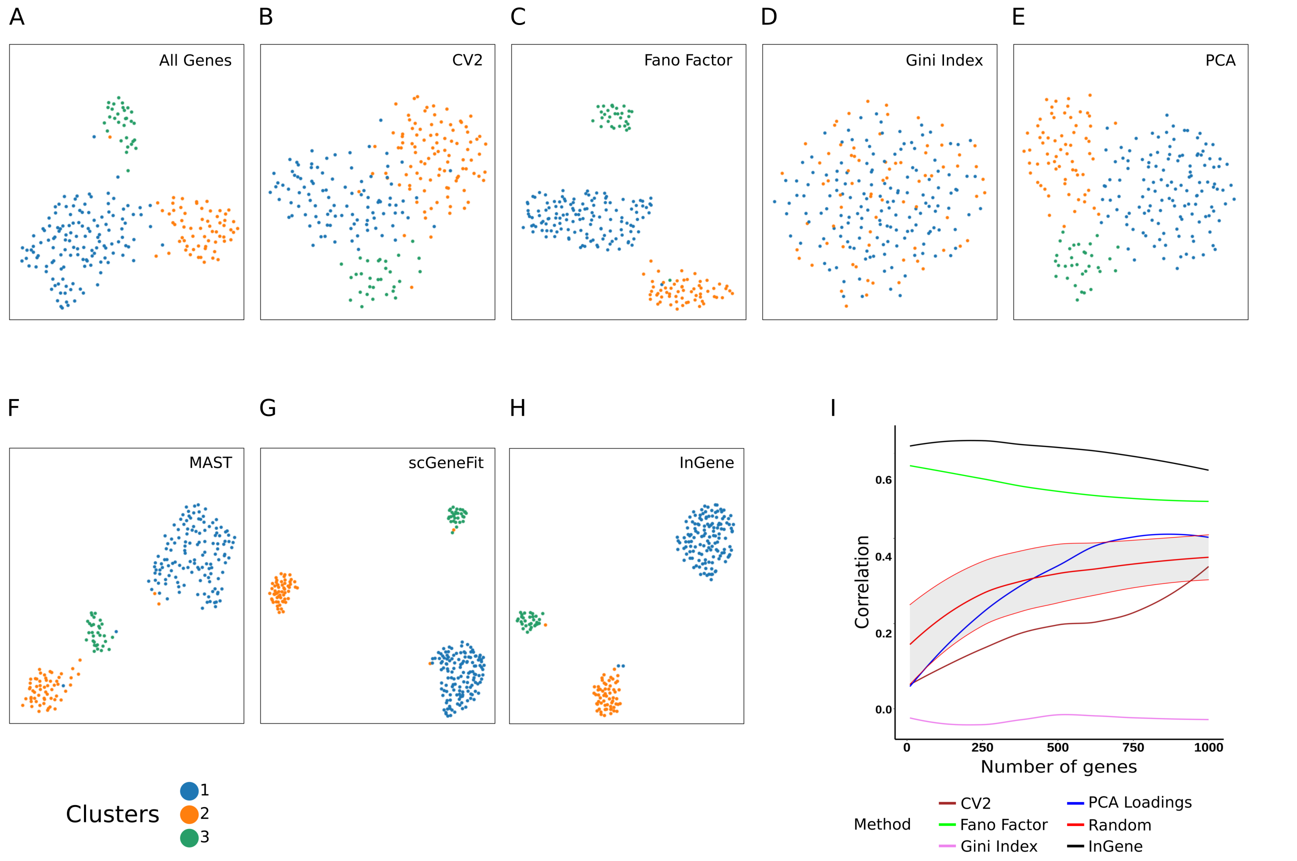


Fig. S 8: Explaining UMAP clusters for single-cell Darmanis dataset with *InGene* **(A)** UMAP constructed with all the genes post-filtering. **(B)** UMAP constructed with top 500 CV2 genes. **(C)** UMAP constructed with top 500 Fano Factor genes. **(D)** UMAP constructed with top 500 Gini Index genes. **(E)** UMAP constructed with top 500 PCA genes. **(F)** UMAP constructed with top 500 MAST genes. **(G)** UMAP constructed with top 500 scGeneFit genes **(H)** UMAP constructed with top 500 *InGene* Factor genes. **(I)** Reconstruction Accuracy (RA) measured for leading 1000 genes for each method. Spearman's rank correlation measures the similarity between high-dimensional and low-dimensional distances. Correlation is calculated with the genes selected in high dimensional space for each method, and compared against randomly selected gene sets of length varying from 10 to 1000 over 20 iterations.

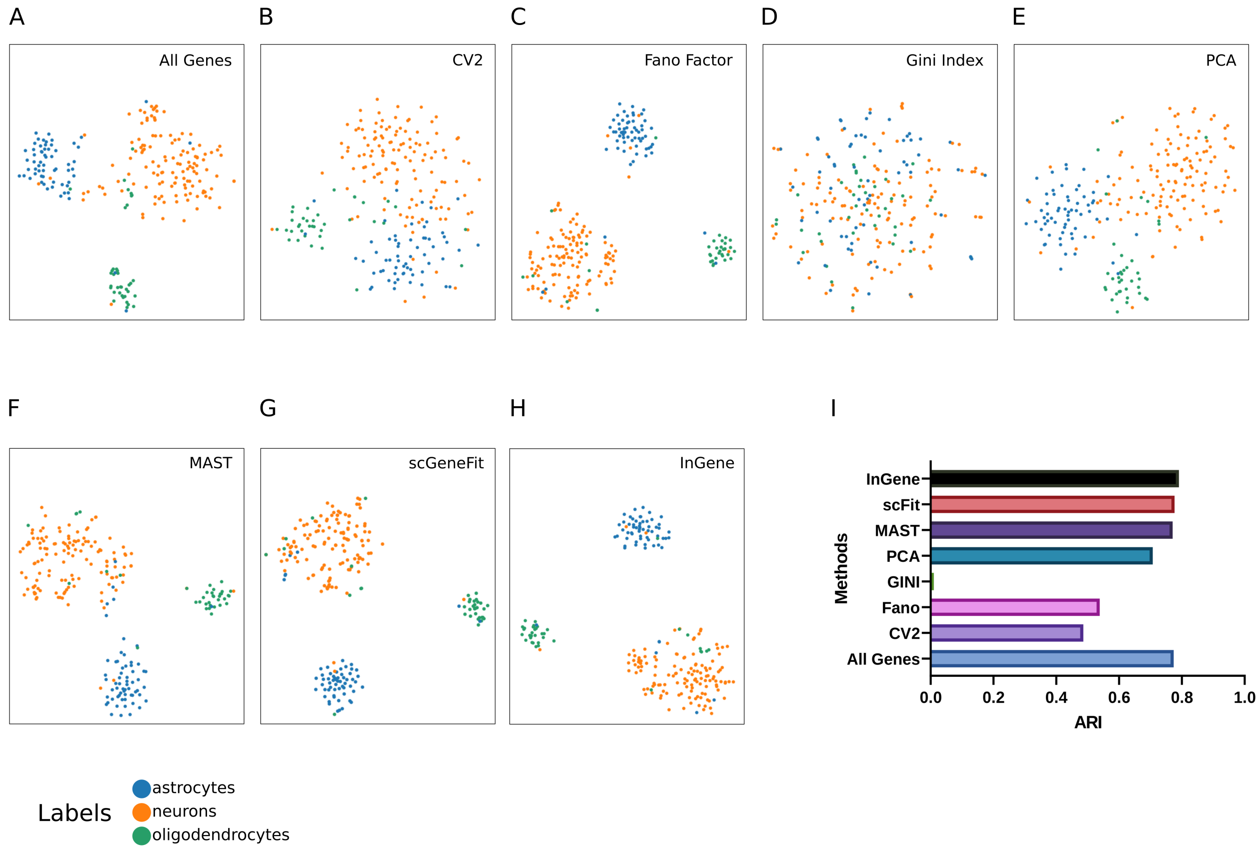


Fig. S 9: Explaining tSNE for single-cell Darmanis dataset with *InGene* **(A)** tSNE constructed with all the genes post-filtering. **(B)** tSNE constructed with top 500 CV2 genes. **(C)** tSNE constructed with top 500 Fano Factor genes. **(D)** tSNE constructed with top 500 Gini Index genes. **(E)** tSNE constructed with top 500 PCA genes. **(F)** tSNE constructed with top 500 MAST genes. **(G)** tSNE constructed with top 500 scGeneFit genes **(H)** tSNE constructed with top 500 *InGene* Factor genes. **(I)** ARI scores for the different methods. The gene set from each methods is used to cluster the dataset, using Leiden algorithm. The cluster labels obtained are then compared with the true labels to obtain the ARI values.

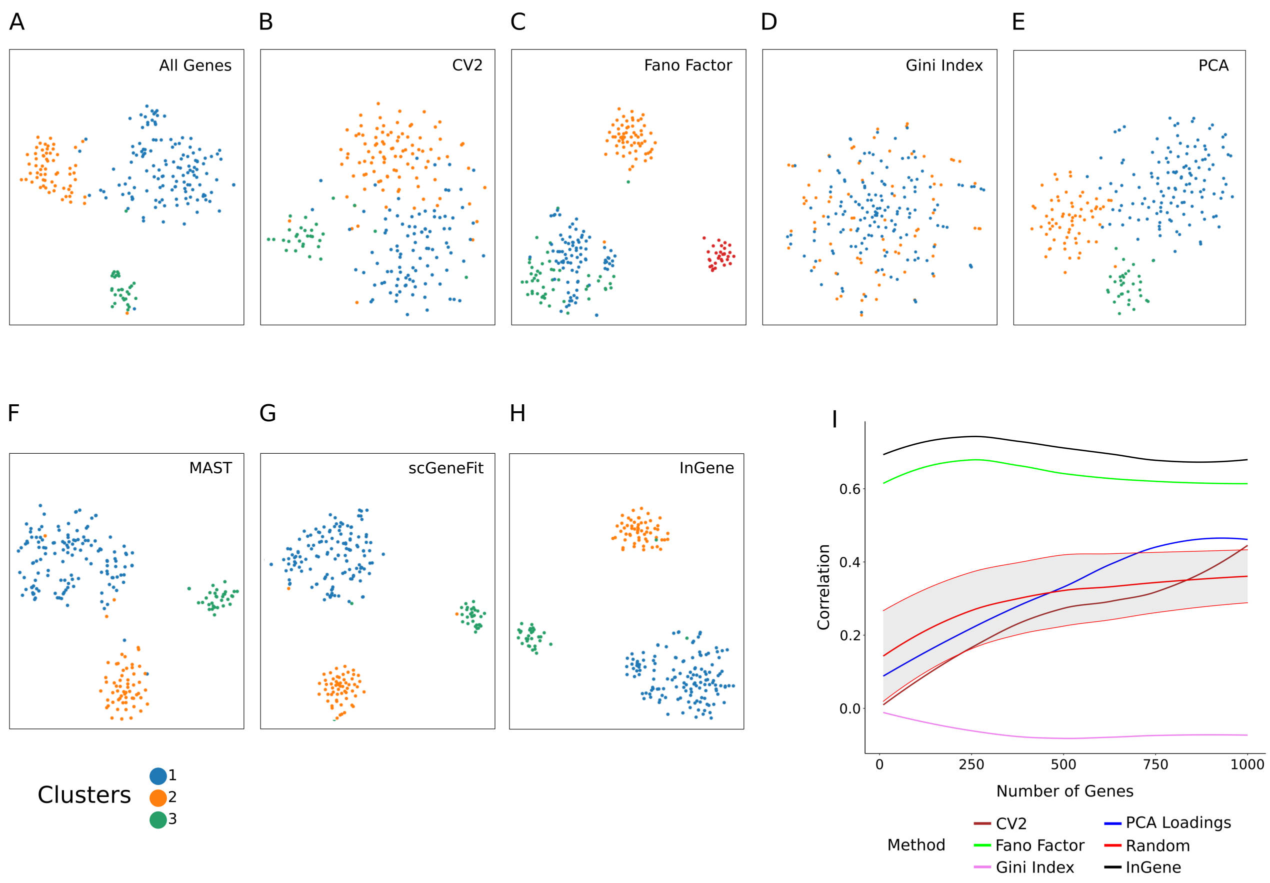


Fig. S 10: Explaining tSNE clusters for single-cell Darmanis dataset with *InGene* **(A)** tSNE constructed with all the genes post-filtering. **(B)** tSNE constructed with top 500 CV2 genes. **(C)** tSNE constructed with top 500 Fano Factor genes. **(D)** tSNE constructed with top 500 Gini Index genes. **(E)** tSNE constructed with top 500 PCA genes. **(F)** tSNE constructed with top 500 MAST genes. **(G)** tSNE constructed with top 500 scGeneFit genes **(H)** tSNE constructed with top 500 *InGene* Factor genes. **(I)** Reconstruction Accuracy (RA) measured for leading 1000 genes for each method. Spearman's rank correlation measures the similarity between high-dimensional and low-dimensional distances. Correlation is calculated with the genes selected in high dimensional space for each method, and compared against randomly selected gene sets of length varying from 10 to 1000 over 20 iterations.

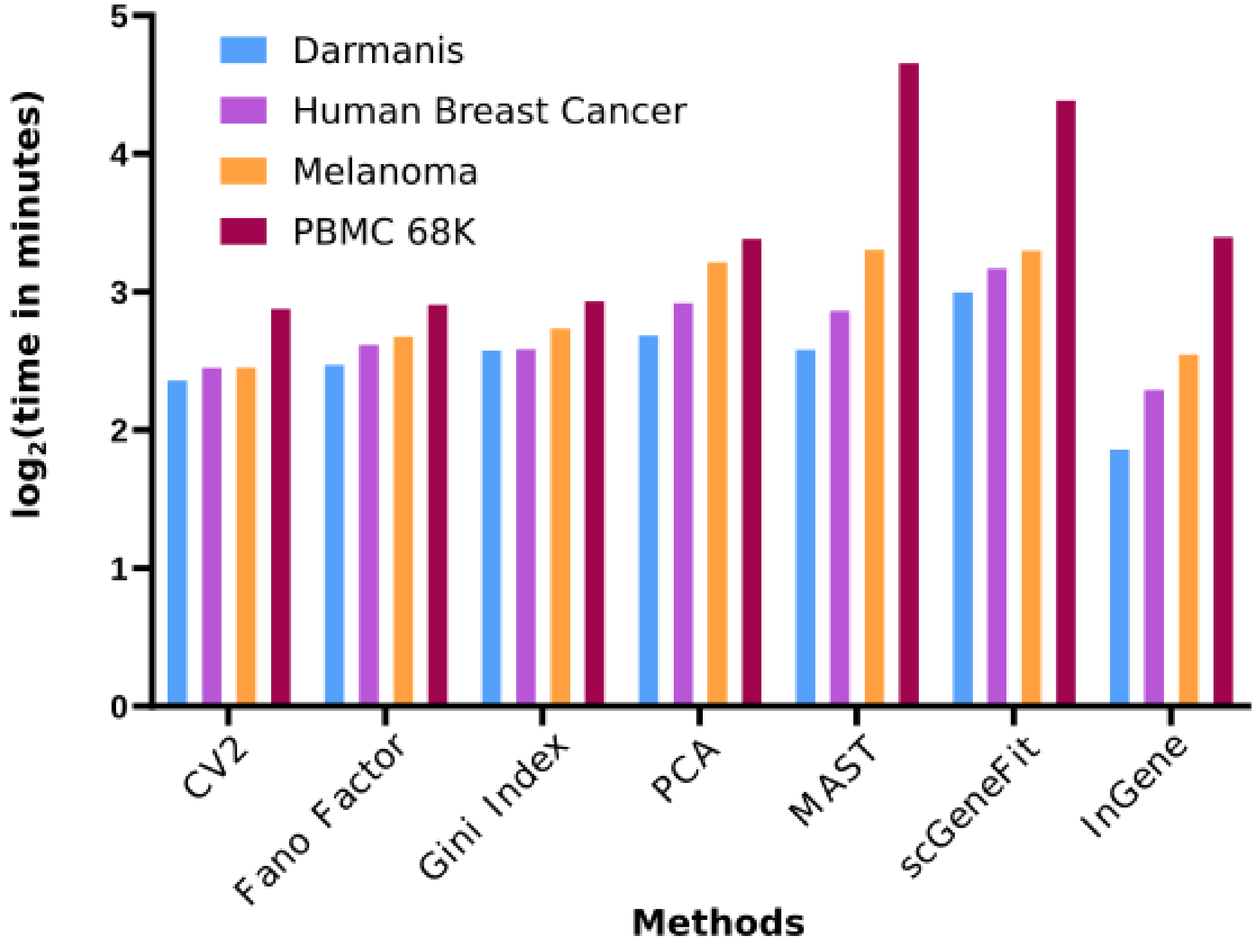


Fig. S 11: *InGene* is scalable. Run time recorded for each method (CV2, Fano Facor, Gini Index, PCA, MAST, scGeneFit, *InGene*) while varying the number of cells from ∼200 to ∼68K.

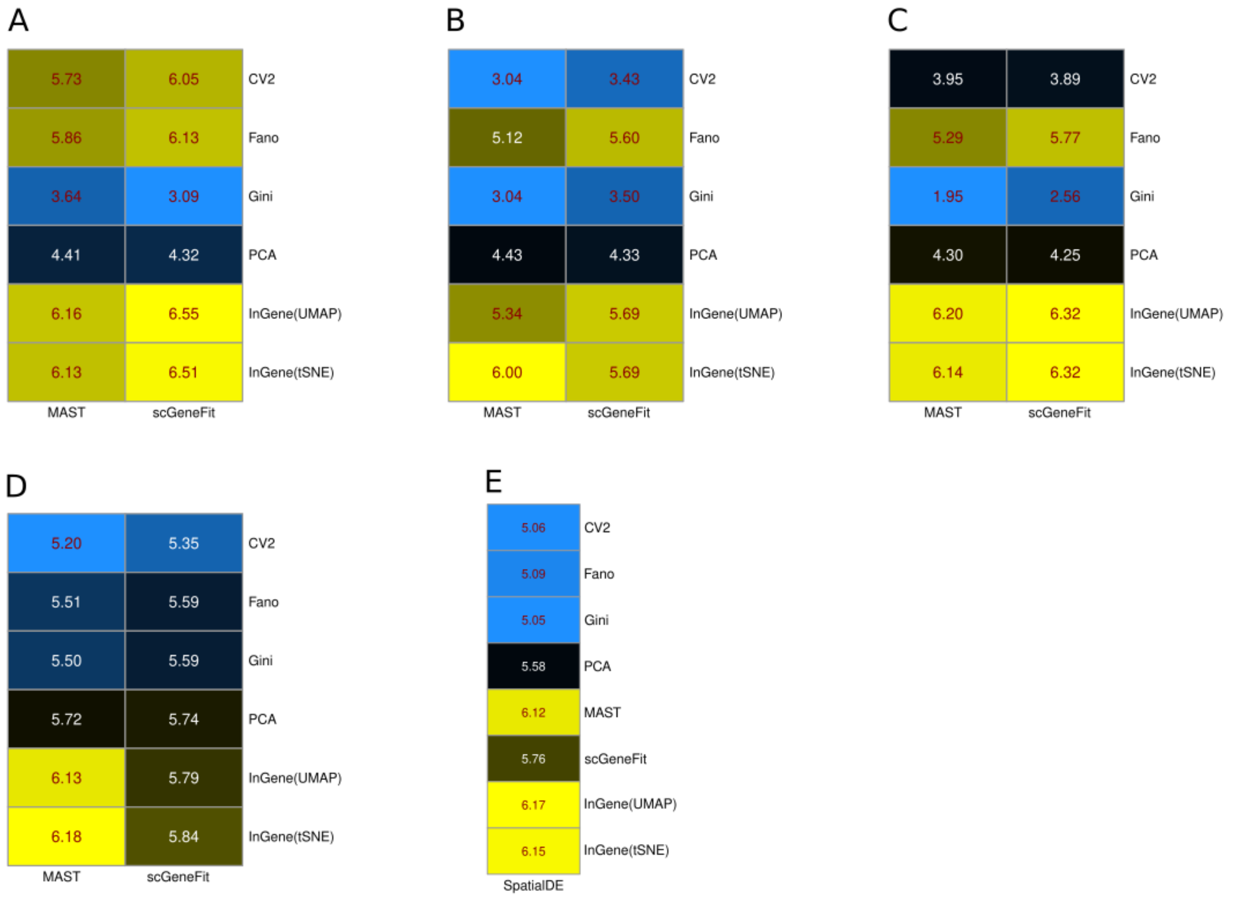


Fig. S 12: *InGene* performs closest to supervised methods. Heatmap (log2-scale) shows that the leading 500 *Ingene* genes has the highest intersection with the supervised methods - MAST, scGeneFit. The comparison is performed for all four datasets. **(A)** Overlap of leading 500 genes from each unsupervised method with those from MAST, scGeneFit, for PBMC 68K dataset. **(B)** Overlap of leading 500 genes from each unsupervised method with those from MAST, scGeneFit, for Melanoma dataset. **(C)** Overlap of leading 500 genes from each unsupervised method with those from MAST, scGeneFit, for Darmanis dataset. **(D)** Overlap of leading 500 genes from each unsupervised method with those from MAST, scGeneFit, for Human Breast Cancer dataset. **(E)** Overlap of leading 500 genes from each method with those from SpatialDE, for Human Breast Cancer dataset.
